# Methylphenidate normalizes aberrant beta oscillations and reduces alpha power during retention in children with ADHD

**DOI:** 10.1101/2020.03.13.990929

**Authors:** C. Mazzetti, N. ter Huurne, J.K. Buitelaar, O. Jensen

**Author notes:** Corresponding author: Cecilia Mazzetti, Donders Institute for Brain, Cognition and Behaviour, Radboud University Nijmegen, P.O. Box 9101, NL-6500 HB Nijmegen, The Netherlands. **Declaration of Interest** Jan K Buitelaar has been in the past 3 years a consultant to / member of advisory board of / and/or speaker for Janssen Cilag BV, Eli Lilly, Medice, Takeda/Shire, Roche, and Servier. He is not an employee of any of these companies, and not a stock shareholder of any of these companies. He has no other financial or material support, including expert testimony, patents, royalties. The remaining authors declare no competing and financial interests.

## Abstract

Attention Deficit-Hyperactivity Disorder (ADHD) has been intensively studied in neurodevelopmental research, with the aim to identify the neural substrates of the disorder. Prior studies have established that brain oscillations in specific frequency ranges associated with attention and motor tasks are altered in ADHD patients as compared to typically developing (TD) peers. We hypothesized that the behavioral improvement following medication in ADHD patients should be accompanied by a normalization in the modulation of such oscillations. We hence implemented a double-blind placebo-controlled crossover design, where boys diagnosed with ADHD underwent behavioral and MEG measurements during a spatial attention task while on and off stimulant medication (methylphenidate, MPH). Results were compared with an age/IQ-matched TD group performing the same task, to assess the effect of MPH on oscillatory activity in the alpha (7 – 13Hz) and beta (15 – 30Hz) bands. We observed that depression of beta band oscillation over motor cortex in preparation to the response in ADHD boys on placebo were significantly lower as compared to the TD group. Importantly MPH resulted in a normalization of the beta depression, which then reached the same levels as in the control subjects. Furthermore, alpha power increased during the preparation interval in the ADHD control group, supposedly reflecting working memory maintenance of the cue information. This increase was significantly reduced in the ADHD group on MPH, reflecting a reduced impact on working memory load. This is the first MEG study showing task related changes in brain oscillations with MPH in children with ADHD.

**Significance statement:** Brain oscillations in the alpha (7-13Hz) and beta (15-30Hz) frequency bands are thought to underly different aspects of attentional processing and their aberrant modulation has been reported in ADHD. Here, we used a child-friendly adaptation of a Posner cueing paradigm to investigate such oscillations in children with and without a diagnosis of ADHD, and further examined the effects of methylphenidate (MPH) in the latter group. We showed that MPH restores aberrant patterns of beta desynchronization and reduces alpha power during retention in the ADHD group, concomitant to an improvement in behavioural performance.

## Introduction

Attention-Deficit Hyperactivity Disorder (ADHD) is a common neurodevelopmental disorder, characterized by a pattern of inattentive and/or hyperactive and impulsive behavior associated with impairments of functioning.

Whereas behavioral therapies have often produced inconsistent results, pharmacological intervention with stimulant medication is widely and effectively used as first line treatment (Faltinsen et al., 2019). The goal of this study was to investigate how the brain network dynamics reflecting the pathophysiology of ADHD responds to administration of MPH in children with ADHD. In particular, we sought to identify psychostimulants’ effects on brain’s oscillatory activity related to attentional processing and motor preparation in children with ADHD. To this end, we examined the influence of MPH on brain oscillatory activity in the alpha (α, 7 – 13Hz) and beta band (β, 13 – 25Hz) in preparation to a motor response to a cued target, in boys with and without ADHD. The effects of MPH on the electrophysiological and behavioral measures of interest were assessed via a double-blind placebo-controlled crossover design within the ADHD group.

Psychostimulants influence midbrain catecholamines levels, particularly norepinephrine and dopamine (Faraone, 2018), by blocking their reuptake in the presynaptic element. In the case of MPH, the biggest effect is thought to be exerted on the increase of synaptic and extra-synaptic dopamine (Tang & Dafny, 2013). Comparable to other kinetic disorders, such as Parkinson’s disease (PD), dopamine disbalance within the fronto-striatal networks observed in ADHD, results into aberrant motor activity and impulsive behavior (Jarczok, Haase, Bluschke, Thiemann, & Bender, 2019), due to both excessive activation and insufficient inhibition along the direct and indirect pathway, respectively (Fernandez-Ruiz et al., 2019).

The therapeutic efficacy of stimulants is widely supported in the control of clinical symptoms in ADHD patients as compared to non-drug treatment options (Coghill, 2019; Faraone & Buitelaar, 2010).

On the other hand, neuroimaging studies have so far provided little insight into the mechanisms of MPH’s action on brain dynamics associated with behavioral improvements. One of the leading hypotheses regarding stimulant’s action on ADHD symptomatology, is that MPH serves to boost distractor suppression, considered a major underpinning of inattention problems in ADHD (Fallon, van der Schaaf, ter Huurne, & Cools, 2016). A previous study linked the improvement in spatial attention due to MPH administration to an increased perception of the saliency of task-relevant information (Niels Ter Huurne et al., 2015). This effect was associated with an improved suppressive gating of irrelevant stimuli to the prefrontal cortex (PFC), due to higher dopaminergic availability. This notion has been corroborated by fMRI studies showing increased BOLD activity within striatal regions concomitant to PFC activation (Fallon et al., 2016). Furthermore, a recent study combined transcranial magnetic stimulation and event-related potentials to investigate the effects of MPH administration on the interaction between cognitive and motor functions (Berger et al., 2018). The study provided evidence for an increased excitability (cortical facilitation) of motor areas concomitant to modulations of attentional processing due to drug intake.

To elucidate the aberrant neural networks underlying motor and attentional symptoms in ADHD, a number of electrophysiological studies have investigated task related modulations of neuronal oscillations in individuals with ADHD. Importantly, aberrant modulations of the sensorimotor μ-rhythm (8 – 12Hz), have been related to motor deficits in adults with ADHD (N ter Huurne, Onnink, Kan, Buitelaar, & Jensen, 2018) and weaker suppressions of β-band activity (13 – 30Hz) has been found in ADHD, prior to response preparation to a cued target (Mazaheri et al., 2014), reflecting weakened appropriate motor planning.

Given their well-established role in attentional selection (Jensen, Bonnefond, & VanRullen, 2012; Jensen & Mazaheri, 2010; Klimesch, 2012), alpha band activity has been another targeted measure of interest in the investigation of neural markers of ADHD. Notably, prior studies have reported a reduced hemispheric lateralization of alpha power in the context of attentional cueing paradigm in children and adults with ADHD compared to controls (Niels Ter Huurne et al., 2013; Vollebregt, Zumer, ter Huurne, Buitelaar, & Jensen, 2016). These findings have been extended in a multimodal imaging context linking aberrant posterior alpha modulations in ADHD to increased frontoparietal connectivity during spatial working memory performance (Lenartowicz, Mazaheri, Jensen, & Loo, 2018). Still, this extensive literature on electrophysiological biomarkers of ADHD lacks a consistent description of the effects of stimulants on such neural indices.

Here, we sought to further elucidate the electrophysiological mechanisms underlying motor and attentional deficits in ADHD, by exploring the effects of MPH administration on sensorimotor beta and posterior alpha oscillations in children with and without ADHD. We hence designed a double-blind placebo-controlled trial to quantify neuromagnetic activity induced by preparation to a motor response in boys under pharmacological treatment for ADHD and typically developing age/IQ matched controls (TD). We speculated that, given that MPH reduces ADHD symptomatology, it would also lead to a normalization of atypical beta and alpha activity patterns, bringing its values closer to the ones observed in the control group.

## Materials and Methods

### Participants

Participants in the study were composed of 27 children diagnosed with ADHD and 27 typically developing (TD) male children. A total of 9 children in the ADHD group withdrew from the experiment after at least one session, due to unwillingness of the parents to continue with the study (N=5), claustrophobic reaction to the MRI scan (N=1), claustrophobic reaction to the MEG acquisition (N=1) and excessive moments during MEG measurements (N=2). One child in the TD group was excluded from the analysis due to a notification of subsequent ADHD diagnosis. As a result, structural and diffusion-weighted MRI scans and MEG data were acquired for a total of 49 (26 TD) and 46 participants (26 TD), respectively. Only the completed datasets (MEG and MRI) were considered for the current study, leaving a total of 18 children diagnosed with ADHD (mean age 10.8±1.0 years) and 26 TD children (10.7±1.2). In the present study, we only focused on electromagnetic results, hence, details on the MRI protocol and data analysis are not further described.

The experiment was conducted in compliance with the declaration of Helsinki and was approved by the local ethics board (CMO region Arnhem-Nijmegen, 2016-2268). All parents gave written informed consent, while children gave oral consent. ADHD children were recruited from different local establishments: the *Karakter Child and Adolescent Psychiatry Centre* in Nijmegen, *Kenniscentrum ADHD and ASS Nijmegen*, *Altrecht Psychiatric Center* in Utrecht and general practitioners’ offices. Children from both groups were recruited via advertisements in local schools and public places, and through the study website.

For both groups, inclusion criteria were: 1) age between 8 and 12 years at the time of the experiment; 2) male; 3) enrolled in primary, not secondary, school; and 4) derived FSIQ measure > 70. For the ADHD diagnosed group, additional inclusion criteria were: 1) a clinical diagnosis of ADHD according to DSM-5 criteria (American Psychiatric Association, 2013); 2) scoring in the clinical range on the ADHD DSM-5 rating scale; 3) being treated with stimulant medication for ADHD (either long or short active formulations), which started at least three months before the inclusion in the study. For both groups, exclusion criteria were: 1) neurological disorders (e.g., epilepsy), currently or in the past; 2) cardiovascular disease, currently or in the past; 3) serious motor or perceptual handicap; 4) standard MRI exclusion criteria (e.g., metal objects/fragments in the body, active implants, brain surgery, claustrophobia). Psychiatric comorbidities were documented, where present, as assessed via the Childhood Behaviour Checklist (CBCL) (Achenbach, 1999), completed by the parents. For the ADHD group, additional comorbidities screening was performed during the psychiatric intake.

### Experimental design

The experiment consisted of two experimental sessions for the TD group, and three sessions for the ADHD group, within a randomized placebo-controlled double-blind crossover design (for a schematic representation of the trial, (see **Figure 1**). Before visiting the institute, parents were asked to fill out the CBCL and ADHD rating scale questionnaires. Parents of the participants in the ADHD group were asked to fill the latter one twice: once referring to the child’s symptomatology pattern while observing him on medication, and once referring to behavior without medication (in this case, the parents were asked to observe their child during the withdrawal period before the experiment). During the first intake-session, children of both groups underwent behavioral testing. This involved a *Line Bisection Test* (LBT), to assess individuals’ spatial biases by asking participants to mark with a pencil the center of a series of horizontal lines (performed both with the left and the right hand) and, if intelligence had not been assessed over the past two years, the *Vocabulary* and *Block Design* subtests of the Wechsler Intelligence Scale (WISC-III), designed for children (Kaufman, 1994; Woolger, 2001); Dutch version in (Kort et al., 2002)), in order to estimate the FSIQ. These subscales have been shown to hold high correlation with the full-scale IQ testing (Herrera-Graf et al., 1996). For the ADHD group, an in-depth intake was conducted by a psychiatrist, where a 30 min interview with parents and son, together with physical examination, were employed to determine the medication dosage to be used during the task. Based on operating procedures followed in prior studies, this led to a medium dosage of 0.3 mg/kg (Linssen et al., 2014). Based on the screening, one of the two standardized dosages was chosen (either 10 or 15mg Methylphenidate immediate release; IR-MPH). Following the behavioral screenings, children of both groups were introduced to the MRI procedure, and were given the opportunity of getting acquainted with the scanner environment by practicing in a so-called ‘*dummy scanner*’, which simulates the environment of a real MRI scanner. If the child was unable to stay still or felt uncomfortable during the dummy scan, the session was cancelled and he was excluded from further participation. Otherwise, the MRI session took place for a duration of ∼30mins.

**Figure 1.**
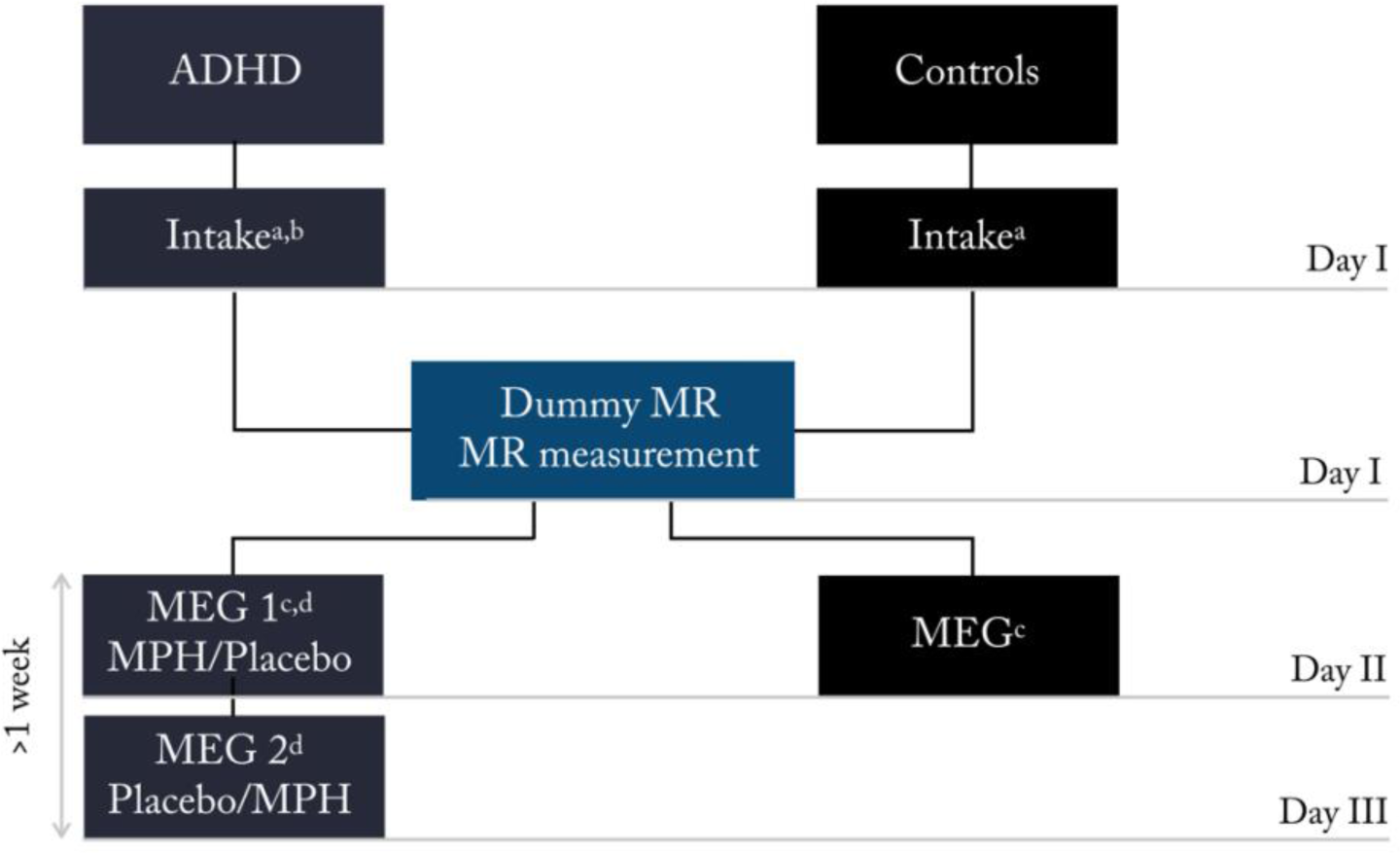
Experimental study design. _a_ Case Report Form and Behavioural test (line bisection task, WISCIII *Vocabulary* and *Block design* subscales, ADHD rating scale, CBCL); _b_ Psychiatric intake (basic medical screening and dosage determination); _c_ Polhemus digitizer; _d_ Medication intake (1 hour prior to the beginning of experimental task); Participants in the ADHD and Controls group visited the laboratory three and two times, respectively. During the first visit (day I) the psychiatric assessment took place, which determined participants’ suitability for the study. The same day, the dummy MR took place, followed by the actual MR scan. The second day (day II) both groups performed the attentional task while electromagnetic activity was recorded with the MEG. The ADHD group performed the task twice (day II and day III), once upon administration of MPH, and once upon administration of a placebo pill. This was done according to a randomized crossover design, half of the participants received the MPH on day II, while the other half on day III.

During the second visit, both groups undertook the MEG testing, followed by 3-D head digitization using an electromagnetic digitizer (*Polhemus*, Colchester, VT). For the TD group, this constituted the last day of testing, while for the ADHD group, two MEG sessions were planned at two different visits, separated by at least one-week interval. Children diagnosed with ADHD performed the MEG task twice under two conditions (MPH and placebo), according to a randomized order and double-blind procedure. Prior to each MEG session, participants were asked to withdraw from their clinically prescribed medication intake for 24 hours, depending on the planned time of the recording session (e.g. if the session was planned in the morning of Monday, the child took his last medication on Sunday morning, abstaining until the experiment). To ensure this, parents agreed on being contacted prior to the experiment to be reminded about the medication withdrawal procedure, to be followed in preparation to the testing day (e.g., in the example above, parents were contacted on Saturday). The 24 hour treatment suspension allowed to control for withdrawal symptoms related to drug administration (rebound effect) (Carlson and Kelly, 2003). MEG testing began one hour after medication intake, allowing to reach on average moderate plasma concentration (*C*_max_) of the drug along the experiment, which progressively increases and reaches its peak around the second hour post-intake (Quinn et al., 2007). After completion of the MEG session, subjects were asked to proceed with their regular treatment using their own stimulant formulation. At the beginning of the experimental session, parents and children were asked to confirm that the medication withdrawal procedure was followed appropriately, else, a new experimental session was planned.

### The attention task

Children from both groups practiced the experimental task prior to the MEG session. The task has its origins in Posner’s cueing paradigm for spatial orienting of attention (Posner and Petersen, 1990) and a related version has previously been applied in pediatric studies investigating attention in ADHD (Vollebregt et al., 2015, 2016). The task was presented as a ‘*save the fish’* game in which sharks were presented on the left and the right side of the screen (**Figure 2**). Experiment was displayed through a PROPixx projector on a screen inside the MEG, holding a resolution of 1920×1080 (aspect ratio: 16:9). Children were asked to maintain fixation on the fish in the middle of the screen. To ensure this, an eye-tracker monitored participants’ eye movements from pre-cue onset until the target presentation. If saccades exceeded the specified limits, set as a boundary box of 100 pixels to the left and to the right and 150 pixels to the top and to the bottom of the center of the screen (fixation point), an error message was presented. This was a video of a talking shark reminding the child to look at the central fish, and an additional trial was presented (allowing us to reject all trials where a correction video was presented). All trials started with of a 500 ms pre-cue period plus the time required to maintain the fixation, as determined by the eye tracker recording. The trial was initiated only when the participant fixated properly (fixation maintained within boundary box). The directional cue (i.e. fish looking to the left or right) was presented for 200 ms, indicating the upcoming position of the target with 80% probability (i.e. the target was in the cued side 80% of the trials and the uncued side in the other 20%). After a preparation interval of 1000 – 1500 ms (jittered), the targets were presented for 300 ms, represented by two tiger sharks at the left and right sides of the screen. The participants had to identify the threatening shark with the open mouth by button press (left- and right-hand button press to indicate respectively left and right target). The child had to respond as fast as possible and within 1100ms. Immediately following the motor response, visual feedback was given, for the duration of 200ms, indicating a correct/incorrect response (happy/dead fish).

**Figure 2.**
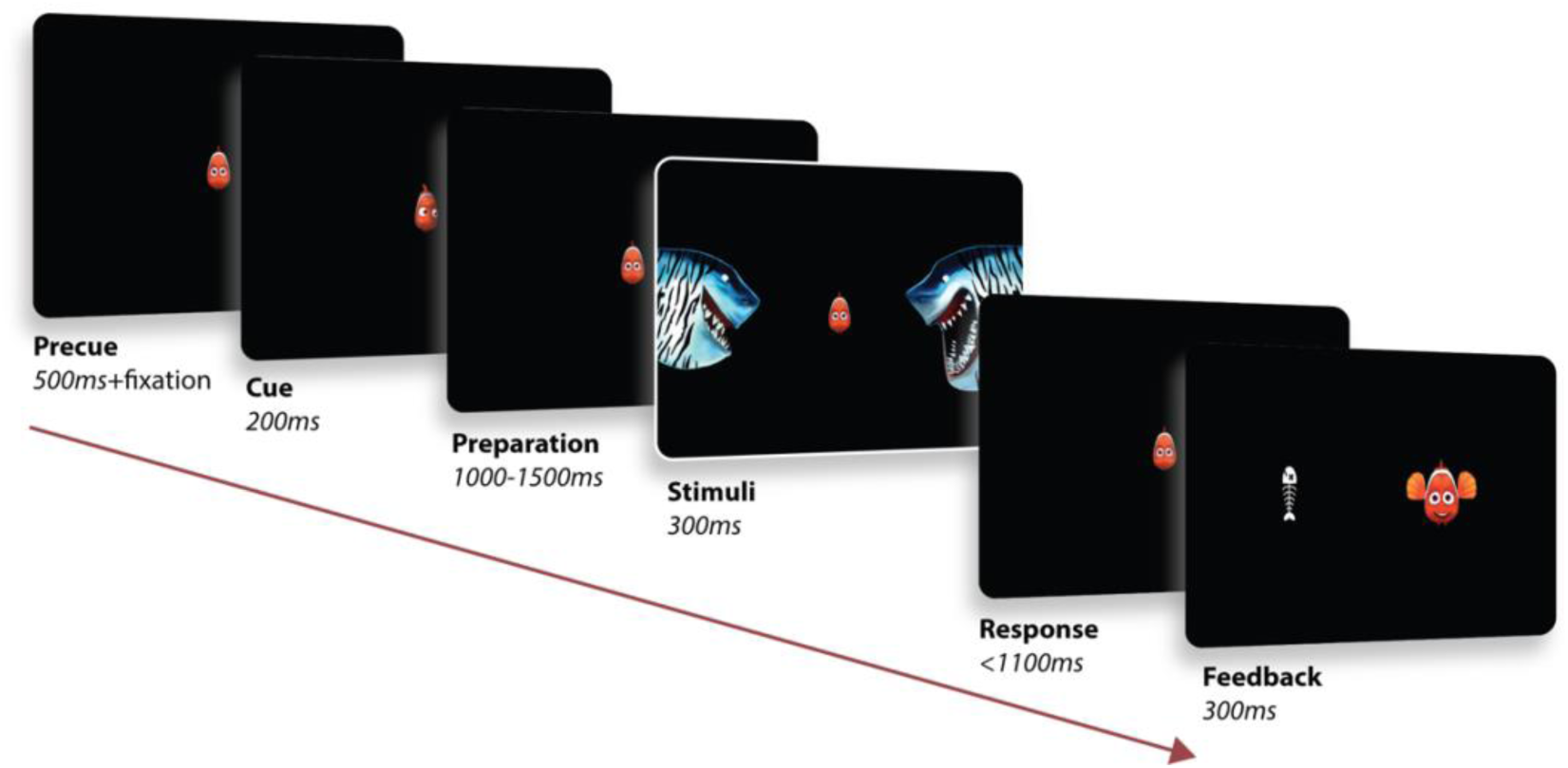
Experimental Paradigm. A child-friendly adaptation of a Posner cueing paradigm for spatial orienting of attention was used. Each trial (370 in total) began with the presentation of a fish in the middle of the screen, serving as fixation cross. An eye tracker ensured that the children kept proper fixation throughout the whole trial (a trial was stopped automatically in case the subject performed a saccade). A cue was then presented for 200ms, represented by the fish looking either at the left or right side of the screen (cue side was equally distributed across trials). After a preparation interval ranging between 1100 – 1500ms (jittered), the stimuli were then presented for 300ms on the two sides of the screen. The child was asked to respond, via button press, indicating the position of the target (shark with an open mouth), while ignoring the distractor on the other side (shark with mouth closed). A positive feedback (happy fish) was then presented if a correct answer was provided within the response interval (1100ms). In case of wrong or no response, a negative feedback was presented (fish bone).

Participants who were unable to perform the task by properly fixating, as monitored through the eye tracker during this practice session as well as during the actual experiment, were excluded from further participation. In case of exclusion from the study, a proper explanation of the reasons was given and children still received a compensation (gift card for a local toy store for the value of €10.00 for each experimental session). The experiment consisted of 360 trials, equally divided in 10 blocks, after which the participant was given the possibility to take a break and/or talk to the parents, if necessary. Note that for the ADHD group the treatment order (methylphenidate versus placebo) for the two MEG sessions was randomized across participants.

### MEG data acquisition

Electromagnetic brain activity was recorded from the participants seated in a CTF 275-sensor whole-head MEG system with axial gradiometers (CTF MEG Systems, VSM MedTech Ltd.). The data were sampled at 1200Hz, following an antialiasing lowpass filter set at 300Hz. Head position was constantly monitored throughout the experiment via online head-localization software. This was done by three head localization coils placed at anatomical fiducials (nasion, left and right ear), allowing, if necessary, readjustment of the participant’s position between blocks.

### MEG data analysis

MEG data analysis was performed using the MATLAB FieldTrip Toolbox (Oostenveld et al., 2011). Initially, the continuous data were segmented in epochs time-locked to the onset of the motor response (−2500 to 1000 ms). A notch filter was applied at 50, 100, 150 Hz to remove line noise, after which the mean was subtracted and the linear trend removed. Trials with incorrect responses according to the cued hemifield were discarded. We then performed a semi-automatic artifacts rejection of trials with MEG sensor jumps and muscle artifacts. Visual inspection was used to further detect and remove trials with eye blinks and systematic saccades which did not exceed the boundary box set by the eye tracker but still produced detectable visual artifacts. ICA was then used to further remove components reflecting eye blinks and heart artifacts.

Prior to the time-frequency analysis of power, we first generated virtual planar gradiometers from spatial derivatives of the axial magnetic components (Bastiaansen and Knösche, 2000), to simplify the interpretation of the topographical mapping of neural activity. Time-frequency representations (TFRs) of power were then computed for each pair of orthogonal planar gradiometers, and power values were then summed for each MEG sensor.

Power analysis was performed using a 600 ms time-window sliding in steps of 50 ms along a time window of interest of −2000 to 200 ms, locked to the onset of the motor response. The resulting data segments were multiplied by a Hanning taper and a fast Fourier transform was applied in the 2 – 40Hz frequency range, in steps of 1.66 Hz. The steps above were also followed for cue-locked epochs, obtained by re-defining the trials according to the onset of the cue, considering the time window of the preceding 1000ms and the following 1000ms.

### Beta power modulation prior to response

In order to estimate patterns of beta modulation indices (β-MI) in preparation to the motor response, we performed a baseline correction (relative change) of response-locked data segments with respect to pre-cue activity per sensor. For each subject, we applied a relative baseline correction with respect to the averaged power in the time window [-300 – 0] ms prior to the cue. The baselined time-frequency representations (TFRs) of power for all subjects were then averaged across conditions and the three groups (TD, ADHD_MPH_, ADHD_Placebo_), in order to identify the sensors of interest to be used in further analyses. To this aim, we selected a cluster of 20 symmetrical pairs of central sensors, displaying the highest beta preparation values (lowest β-MI) in the 1000ms time-window preceding the onset of the motor response. The same sensors were then used for the estimation of mean beta modulation indices (PI(β)) for each subject, by averaging MI(β)s across the time window of interest.

Additionally, we quantified the individual variability in interhemispheric modulation of beta power (HMI(β)) by computing, for each subject, average PI(β)s over left and right ROIs.

### Beta power lateralization prior to response

We then considered individual TFRs, separately for *attend* left and *attend* right trials. A Lateralized Modulation Index (LMI) was computed for each sensor *k* and over all time points *t* belonging to the time window of interest −1000 – 0 ms, according to the formula:

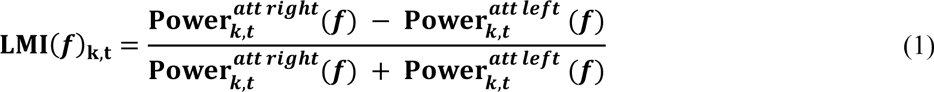

Where *Power*(*f*)_*k,t att left*_ represents the power at a given frequency *f* in the condition ‘attend left’ and *Power*(*f*)_*k,t* att right_ is the power of the same frequency in the condition ‘attend right’. As a result, positive (or negative) LMI values, at a given sensor *k* and given timepoint *t*, indicate higher power at a given frequency *f* when the subject prepared to a motor response to a cued right (or left) target (i.e. right or left finger). Two clusters of sensors were derived, by selecting the twenty symmetrical occipito-parietal sensors (i.e. ten pairs of sensors) showing the highest interhemispheric difference in beta lateralized modulation indices (LI(β)), when considering the grandaverage over all conditions (see **Figure 10**) averaged over the previously defined time window of interest (−1000 – 0 ms). These clusters constituted the regions of interests (ROIs) on which subsequent analysis of lateralized modulation was focused. In order to derive an index of individual interhemispheric modulation, we calculated Beta Lateralization Indices (LI(β)) per participant, according to:

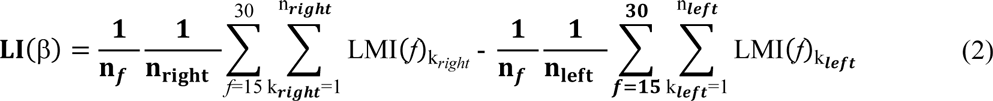

Where LI is averaged in the interval of β (15-30Hz) and *k_left_* and *k_right_* denote sensors belonging to the left and right clusters of selected sensors, respectively. Since LMI(β) values were obtained by contrasting beta power in right versus left attention trials (see Eq.1), left hemisphere LMI(β) are represented by negative values, and right hemisphere LMI(β) by positive values. Consequently, higher LMI(β) indicated higher alpha lateralization for a given subject (i.e., higher interhemispheric difference in absolute beta modulation).

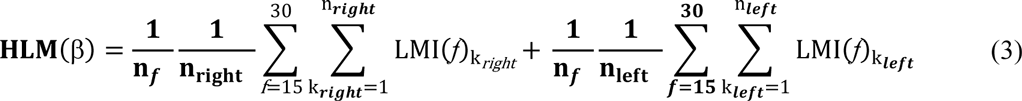

Hemispheric Lateralized Modulation index of beta power (HLM(β)) were computed as follows:

As in the previous formula, *f* is averaged over the β (15-30Hz), and, equivalent to the terms included in Eq.(2), left hemisphere LMI(β) are negative while right hemisphere LMI(β) are positive. By summing their values, the HLM(β) is positive if lateralized beta modulation is higher in the right hemisphere, and negative if higher in the left hemisphere.

### Cue-locked analysis (α-band activity)

For attention-related activity, epochs were redefined to be time-locked the onset of the cue, considering a time window of interest between −2500 and 1500ms. Consistent with the procedure followed for the analysis of beta power, we derived virtual planar gradiometers for the computations of TFRs. Time-frequency analysis of power was performed using a 600ms sliding time-window in steps of 50ms, over the whole 4s epoch. The resulting data segments were multiplied by a Hanning taper and a fast Fourier transform was applied in the 2 – 40Hz frequency range, in steps of 1.66 Hz.

### Alpha power modulation following the spatial cue

Comparable to the analysis steps followed for the beta range, we identified oscillatory patterns of α-band (*f*=7-13Hz) activity in the preparation interval (i.e. following the spatial cue and preceding the stimulus onset) by computing relative power change and evaluating the modulation in the 400 – 1200ms interval with respect to the onset of the cue.

### Alpha power lateralization following the spatial cue

In order to quantify individual interhemispheric variability of alpha modulation following a spatial cue, while preparing for the response to the target (preparation interval), we considered the TFRs of power locked to the onset of the cue. We then considered individual TFRs, separately for *attend* left and *attend* right trials. A Lateralized Modulation Index (LMI) was then computed for each sensor *k* and over all time points *t* belonging to the time window of interest 400 – 1200ms, according to Eq.(1). We then selected two clusters of symmetrical sensors showing the highest paired interhemispheric difference in alpha power 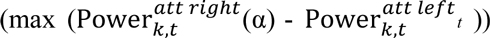 from occipital and posterior areas, when considering the grand average of the three conditions (groups). These clusters constituted the regions of interests (ROIs) on which subsequent analysis of lateralized modulation was focused.

Measures of individual interhemispheric alpha modulation were derived by computing Lateralization Indices in the alpha band (LI(α)) per participant, according to Eq(2).

Next, Eq(3) was then applied to the alpha frequency band to compute Hemispheric Lateralized Modulation index of α power (HLM(α)): as for the beta band, left hemisphere LMI(α) are negative while right hemisphere LMI(α) are positive, giving a positive HLM(α) for a higher α lateralized modulation in the right hemisphere, and negative if higher in the left hemisphere.

### Individual alpha peaks

To identify individual alpha peaks (αP), we first selected a set of sensors of interest comprising parietal and occipital regions. We then computed power spectra over 1s time-window post-cue (200ms-1200ms), applying zero padding to 10s and multiplying the resulting segments with a Hanning taper. Power between 2-20Hz was calculated using a Fast Fourier Transform (FFT). Peaks were included when in the range between 7 – 14Hz, and, in case of multiple peaks, the peak associated with highest power was chosen. Alpha peak power modulation (MI(αP)), peak retention index RI(αP), lateralized modulation index (LMI(αP)) and lateralization index (LI(αP)) were calculated according to the same methods previously described for the alpha frequency band (Eq. 1,2 and 3).

### Behavioral and electrophysiological indices related to the task are not affected by drug nor session order

Given the cross-over design, we controlled for the presence of crossover (*drug condition*: MPH vs Placebo) and sequence effects (MEG *session order*). These factors can be of potential confound whereas familiarization with the task led to overall better performance on the second experimental session. In order to assess whether the interaction of drug condition and session order influenced the reported measures of interest, we implemented a general linear model where *drug condition* and *session order* were defined as regressors for the predictors of the main behavioral and electrophysiological indices: Inverse efficiency scores (IES), cueing effect indices (CEI), beta preparation (PI(β)), beta lateralization (LI(β)), alpha increase (RI(α)), alpha lateralization (LI(α)). No significant interaction effects of Drug*session order arose from any of the models specified except for the alpha lateralization indices LI(α) (see **Table 3**). From a statistical point of view, this meant that any further investigation of LI(α) and its association with other task variables, had to control for the variance explained by the session order. For the other comparisons, this confirmed that our results were not affected by the sequence of experimental sessions.

**Table 1.** Clinical features of the ADHD sample

| ADHD Clinical features | prevalence<br>(N=18) | % |
| --- | --- | --- |
| **Treatment medication** |  |  |
| Concerta | 5 | 27.8 |
| Ritalin | 5 | 27.8 |
| Methylphenidate | 5 | 27.8 |
| Medikinet | 3 | 16.7 |
| Equasym | 2 | 11.1 |
| **Comorbidity** |  |  |
| ODD | 1 | 5.6 |
| PDD-NOS | 1 | 5.6 |
| Learning disabilities NOS | 1 | 5.6 |
| AD-NOS | 1 | 5.6 |
| ASS | 1 | 5.6 |
| Tic Disorder | 2 | 11.1 |
| Constitutional Eczema | 1 | 5.6 |
| **Dosage MPH-IR administered** |  |  |
| 15mg | 13 | 72.2 |
| 10mg | 5 | 27.8 |
| **ADHD Subtype** |  |  |
| inattentive | 1 | 5.6 |
| combined | 4 | 22.2 |
| impulsive | 1 | 5.6 |

**Table 2.** Demographics and summary of statistics between groups. *For a complete overview of comparisons of ADHD scores within ADHD group and between groups see **Figure 12**.

| Descriptive Characteristics | TD (N=26) | ADHD Placebo (N=18) | Statistics |
| --- | --- | --- | --- |
| Age | 10.7 (1.2) | 10.8 (1.0) | n.s. 1145 |
| Estimated IQ | 126.0 (20.1) | 111.1 (34.8) | n.s. 1146 |
| Line Bisection Task | 0.51 (0.02) | 0.50 (0.01) | n.s. 1147 |
| Handedness (%) |  |  |  |
| left | 4 | 7.1 | n.s. 1148 |
| right | 96 | 92.9 | n.s. 1149 |
| ADHD Scores* |  |  | 1150 |
| ADHD Inattention | 5.85 (4.10) | 20.33 (3.34) | p<.001 1151 |
| ADHD Hyperactivity | 1.60 (1.09) | 12.22 (3.45) | p<.001 1152 |
| ADHD Impulsivity | 1.32 (1.65) | 6.88 (1.81) | p<.001 1153 |

**Table 3.**
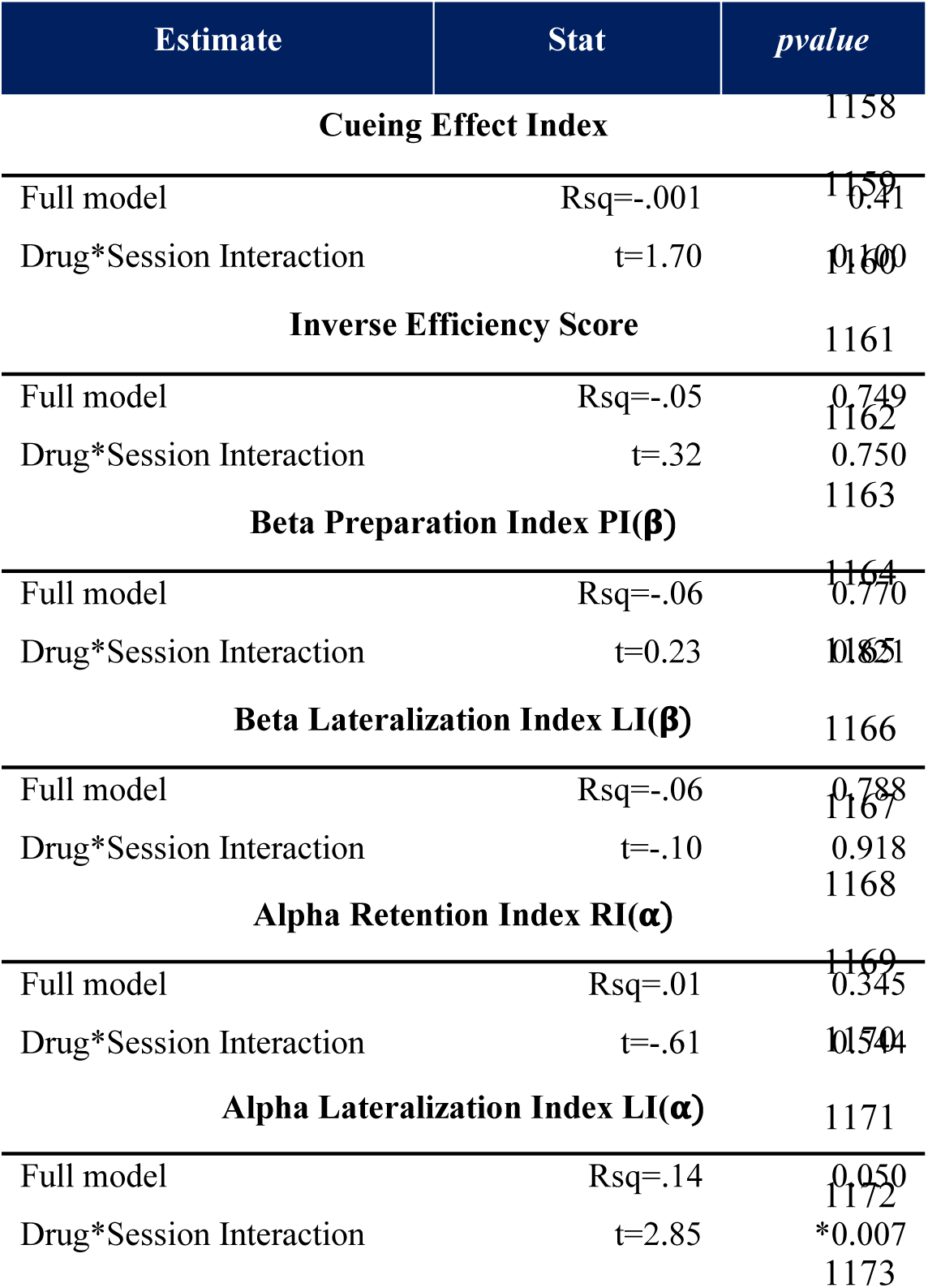
Crossover and sequence effects for main variables of interest in the task

| Estimate | Stat | *pvalue* |
| --- | --- | --- |
| **Cueing Effect Index** |  | 1158 |
| Full model | Rsq=-.001 | 1159<br>0.41 |
| Drug*Session Interaction | t=1.70 | 1160<br>0.10 |
| **Inverse Efficiency Score** |  | 1161 |
| Full model | Rsq=-.05 | 1162<br>0.749 |
| Drug*Session Interaction | t=.32 | 1163<br>0.750 |
| **Beta Preparation Index PI($\beta$)** |  | 1164 |
| Full model | Rsq=-.06 | 1165<br>0.770 |
| Drug*Session Interaction | t=0.23 | 1166<br>0.821 |
| **Beta Lateralization Index LI($\beta$)** |  | 1166 |
| Full model | Rsq=-.06 | 1167<br>0.788 |
| Drug*Session Interaction | t=-.10 | 1168<br>0.918 |
| **Alpha Retention Index RI($\alpha$)** |  | 1169 |
| Full model | Rsq=.01 | 1170<br>0.345 |
| Drug*Session Interaction | t=-.61 | 1171<br>0.541 |
| **Alpha Lateralization Index LI($\alpha$)** |  | 1171 |
| Full model | Rsq=.14 | 1172<br>0.050 |
| Drug*Session Interaction | t=2.85 | 1173<br>*0.007 |

**Table 4.** Summary of Alpha peaks statistics

| Comparison | Statistics |  |
| --- | --- | --- |
|  | Estimate | pvalue |
| **Alpha Peak Retention Index RI(αP)** |  |  |
| MPH vs Plac | t(17)= 2.12 | 0.048 |
| MPH vs TD | t(42)= 1.01 | 0.319 |
| Plac vs TD | t(42)=-0.42 | 0.674 |
| **Alpha Peak Lateralization Index LI(αP)** |  |  |
| MPH vs Plac | t(17)= 0.41 | 0.688 |
| MPH vs TD | t(42)= 0.65 | 0.515 |
| Plac vs TD | t(42)= 0.23 | 0.813 |
| **RI(αP) corr IES** |  |  |
| MPH | r=-0.20 | 0.428 |
| Plac | r=0.003 | 0.990 |
| TD | r=-0.16 | 0.432 |
| **ΔIES ~Δ-RI(αP) \*Dosage** |  |  |
| ΔRI(αP) | t =-1.35 | 0.196 |
| Dosage | t =-1.62 | 0.126 |
| ΔRI(αP) *Dosage | t = 1.41 | 0.178 |
| **LBT corr LI(αP)** |  |  |
| MPH | r= 0.08 | 0.750 |
| Plac | r=-0.06 | 0.756 |
| TD | r= 0.44 | 0.063 |
| **HLM(αP)** |  |  |
| MPH vs Plac | t(17)= 0.34 | 0.736 |
| MPH vs TD | t(42)=-0.87 | 0.385 |
| Plac vs TD | t(42)=-1.50 | 0.142 |
| **LBT corr HLM(αP)** |  |  |
| MPH | r= 0.11 | 0.657 |
| Plac | r= 0.12 | 0.638 |
| TD | r=-0.08 | 0.674 |

## Results

In the current study, we acquired structural MRI and electrophysiological MEG data from 44 male children (26 typically-developing (TD) children and 18 children diagnosed with ADHD). Participants’ attentional performance was tested in child-friendly spatial cueing paradigm (see **Figure 2**) and the task-modulated brain activity was then characterized.

### MPH improves behavioural task performance in the ADHD group

To assess whether behavioural performance in the task improved with medication intake, we computed an Inverse Efficiency Scores (IES) per subject, by taking into account both accuracy (Acc; defined as hits divided by misses) and reaction times (RT) according to:

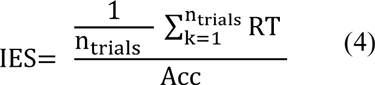

This measure is effective in assessing behavioural performance by balancing the speed-accuracy trade-off (Vandierendonck, 2017). Importantly, MPH administration led to a significant improvement in task performance (t_(17)_=2.49, *p*=.023) in the ADHD group, as reflected by lower mean IESs (**Figure 3**). No significant difference in mean IES was instead found between TD and ADHD: the TD group did not perform better when compared to the ADHD group in the Placebo (t_(42)_= −.22, *p*=.827) nor in the MPH condition (t_(42)_= .77, *p*=.445).

**Figure 3.**
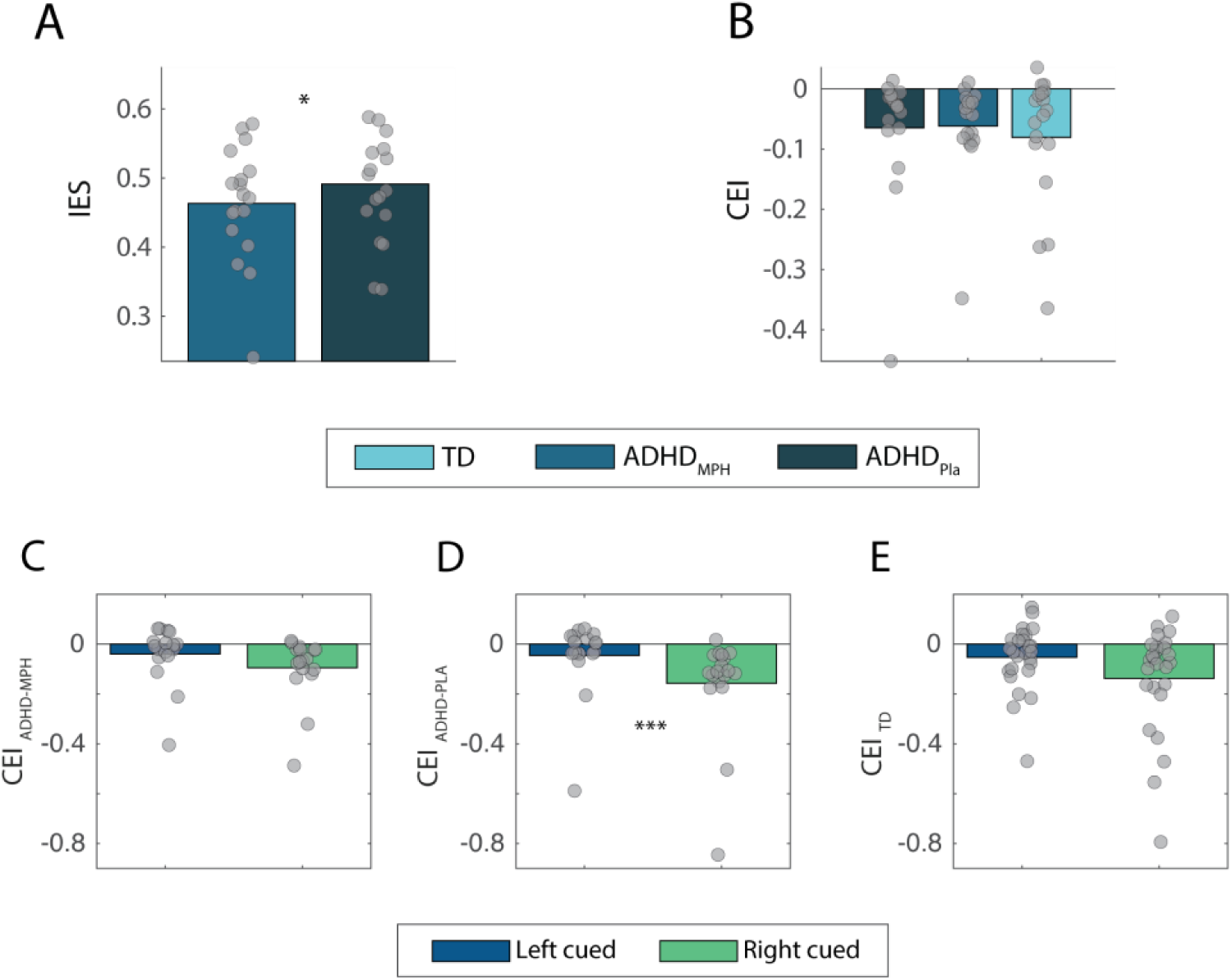
MPH administration leads to improved behavioural performance. (A) Bars show inverse efficiency scores (IESs: median RTs divided by accuracy) averaged for the ADHD groups in the MPH or Placebo condition. Individual IES scores are superimposed and represented by grey dots. A significant difference in IES between MPH (blue) and placebo administration (dark blue), demonstrated that MPH administration led to improved task performance. (B) Cueing Effect Indices (CEI) averaged over the three groups. No difference in mean CEI was observed between conditions. (C) perceptual hemifield bias was quantified by separately computing CEI for left and right cued conditions in all three groups. While no inter-hemifield difference was found for the ADHD_MPH_ (C) nor the TD (E) group, a significantly higher cue benefit (i.e., smaller CEI) was found for the ADHD_PLA_ group (D).

To assess whether subjects benefited from the presence of the cue, we computed IES separately for validly and invalidly cued trials and compared them within groups: as a result, only the TD group showed a significant improvement in performance for valid as compared to invalid trials (t_(25)_= −3.04, *p*=.015, *corrected*), while ADHD participants in both the Placebo and MPH condition did not significantly benefit from cue direction (t_(17)_=−1.70, *p*=.330 and t_(16)_=−2.11, *p*=.150, *corrected*).

In line with previous studies investigating attentional performance in clinical samples (ter Huurne et al., 2013; Vollebregt et al., 2015), we also computed a Cueing Effect Index (CEI), defined as IES for valid trials minus IES for invalid trials, normalized by their sum. As a result, the smaller (more negative) the CEI, the faster were the responses to cued targets (higher benefit from the cue).

No difference in mean CEI was observed between conditions (ADHD_MPH_ vs ADHD_PLA_: t_(17)_=.29, *p*= .775; ADHD_MPH_ vs TD : t_(42)_=−.07, *p*= .942; ADHD_PLA_ vs TD : t_(42)_=−.28, *p*= .779).

### Perceptual and motor interhemispheric biases

For all subjects, we used a Line Bisection Task to evaluate the existence of motor and perceptual biases (see *Materials and Methods*). To avoid confounding effects from hand dominance, we asked participants to perform the task twice, once with the left and once with the right hand. Average values were then computed and evaluated within and between groups, to identify, if any, pseudoneglect biases within and between groups. No average unilateral spatial bias was found in neither of the two groups (neglect being indicated where mark position exceeded 10% of average line half-length (McIntosh et al., 2017)) and no significant difference in directionality was found between groups (t_(42)_=−.73, *p*=.470).

Perceptual hemifield biases for each experimental group were assessed by comparing the cue benefit between left and right trials (CEI). A significantly higher CEI for right cued trials was found for the ADHD group on Placebo (t_(17)_=−5.89, *p*=5.36×10_-5,_ *corrected*, **Figure 3D**), confirming a right hemifield preference for ADHD children reported in previous literature (Wagner et al., 2015). The ADHD_MPH_ and TD group did not show this perceptual bias (t_(17)_=−2.36, *p*=.093 and t_(25)_=−2.22, *p*=.106, *corrected*, **Figure 3C,E**).

A two-way repeated measures ANOVA was then implemented to assess the main effects of *drug condition* (MPH vs Pla) and *hemifield* (left vs right cues) and their interaction on the CEI in the ADHD group. A main effect for *hemifield* was found (F_1,64_= 9.34, *p*=.003), but not for *drug condition* (F_1,64_= 0.01, *p*=.930). The interaction *drug condition* * *hemifield* yielded no significant effect (F_1,64_=0.84, *p*= .362). In conclusion, the right bias found in ADHD_Pla_ group was not significantly reduced with MPH.

### Methylphenidate normalizes aberrant beta depression patterns in ADHD children

Participants in the ADHD group were tested on the task twice, namely while methylphenidate (10mg or 15mg) or placebo was administered, within a double-blind crossover design. After appropriate preprocessing and artifact rejection, time-frequency representations of power (TFRs) were computed for all valid correct trials locked to the onset of the motor response (i.e. button press, see *Materials and Methods* for details). Trials were baseline-corrected using the spectral power in the −300 – 0 ms pre-cue interval. Our first aim was to identify the sensors and time-ranges associated with the motor preparation prior to the button press. The topographical representation of mean MI(β)s in the TF spectrum of interest (15 – 30Hz, −1000 < t < 0 ms) across all conditions is presented in **Figure 4A**: a clear depression in beta band power (β-MI) was present at central sensors in the interval preceding the button-press when averaging over trials, conditions and subjects. For further analysis, we selected a cluster of 20 symmetrical pairs of central sensors showing the highest β-MI (i.e. highest beta power depression; **Figure 4A**) along the −1 – 0 s time-interval with respect to the button press (**Figure 4B**).

**Figure 4.**
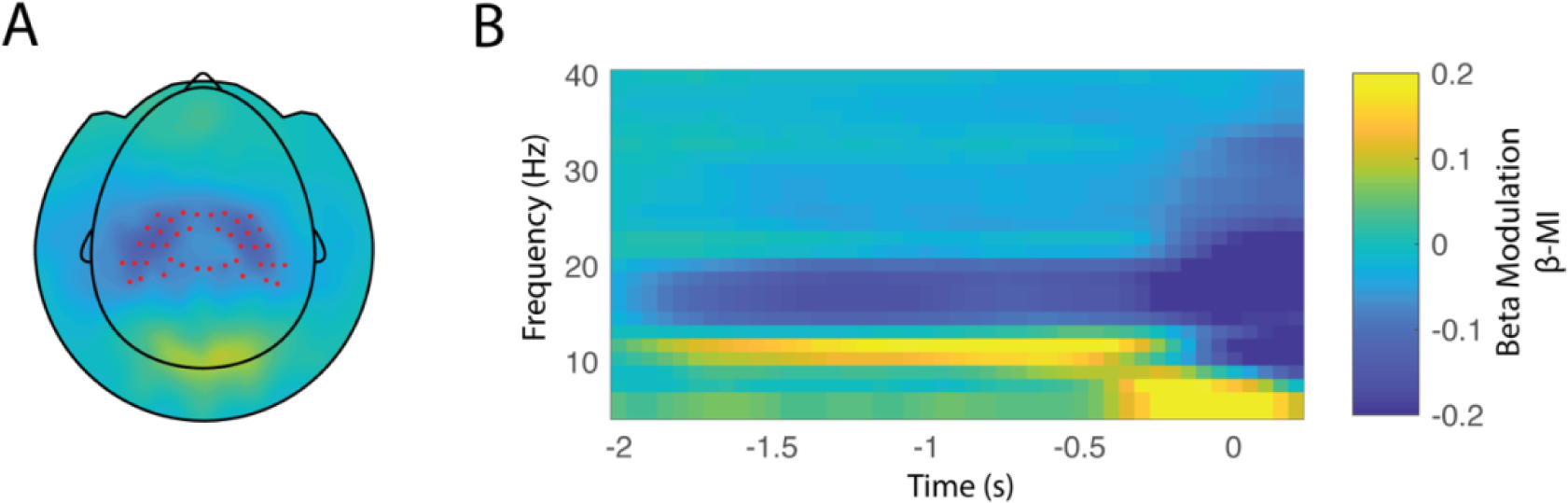
Grandaverage beta modulation locked to response to the cued target. (A) Topographic representation of the mean beta modulation index (15-30Hz) across all conditions in the time window of interest ([-1 – 0]s) prior to the response. A clear depression in the power of beta oscillations is visible at central sensors. Pairs of symmetric sensors displaying the highest beta depression values, are marked with red dots and will be used as sensors-of-interest (B) Time-frequency representation (TFR) of frequency power modulation averaged across conditions. Power values shown were calculated across the cluster of sensors highlighted in (A) and locked to the onset of the motor response to the target.

**Figure 5** illustrates the respective topographical plots and TFRs from the sensors and time-window of interest, for the TD, MPH and Placebo groups (**Figure 5**). In all groups, we observed a depression of beta power reflecting the preparation for the button press (**Figure 6A**). To compare the beta power depression (PI(β)) between conditions, we averaged MI(β)s in the time-frequency representations and sensors of interest for the three conditions (**Figure 6B**). Independent sample t-test was employed to compare the PI(β)s between TD and ADHD children. We first observed that beta depression was significantly higher in the TD compared to ADHD group in the Placebo condition (t_(42)_=−2.52, *p*=.016), but not compared to the MPH condition. Importantly, a paired t-test revealed a significant difference within the ADHD group as a result of the treatment: more pronounced beta depression was found for ADHD group administered with MPH compared to the Placebo condition (t_(17)_=2.36, *p*=.028). In sum these findings show that preparatory beta depression is reduced in children with ADHD, and that administration of MPH serves to restore beta oscillations during preparation to response.

**Figure 5.**
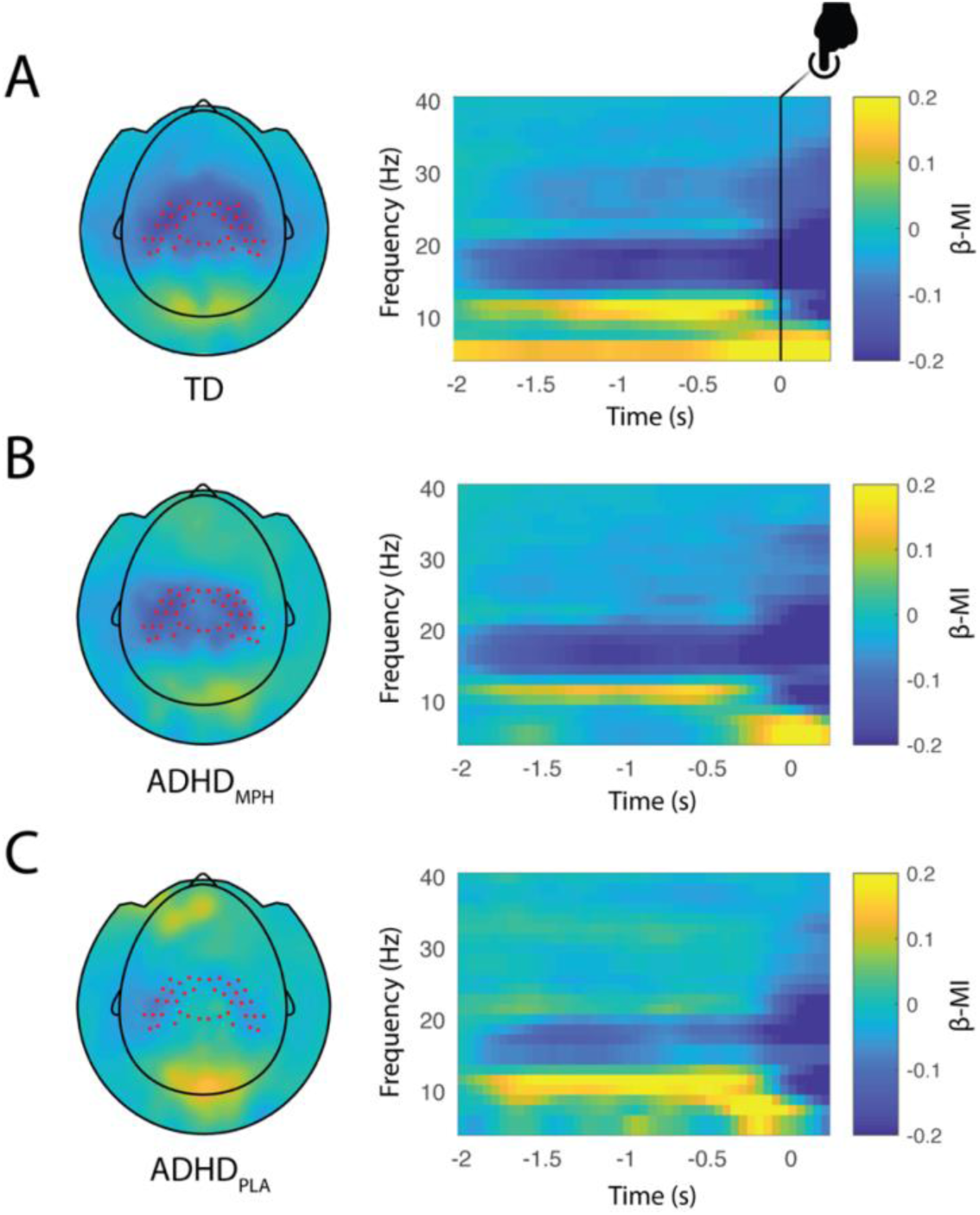
Beta modulation indices in the three conditions. Topographic plot (left) and respective time frequency representations (TFRs) (right panel) of power modulation for the typically developing group (TD) (A), ADHD_MPH_ group (B) and ADHD_PLA_ group (C). Red dots superimposed on the topographies denote sensors of interest as defined in Figure 4A. Notably, beta preparation is stronger in the TD group, while progressively decreases in the ADHD_MPH_ group, being weakest in the ADHD_PLA_ group.

**Figure 6.**
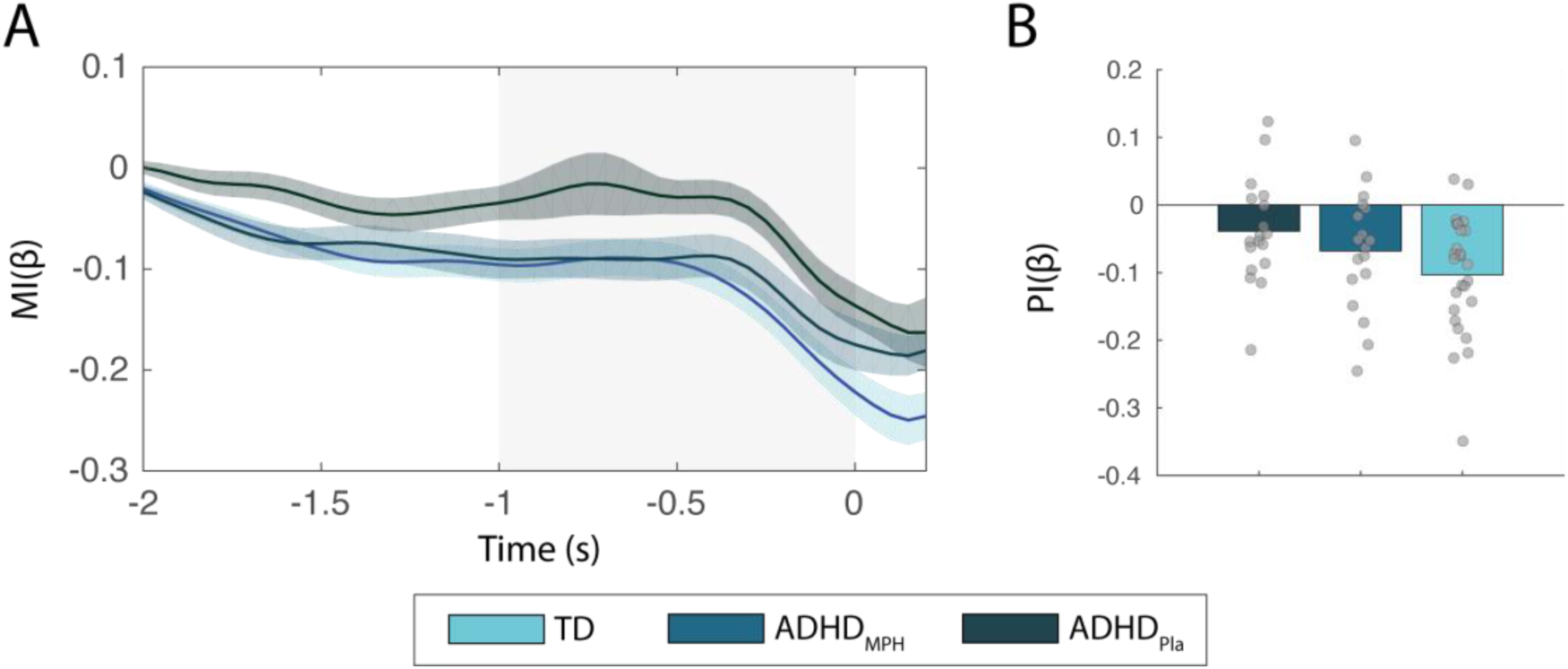
Evolution of MI(β)s over time and mean beta preparation index (PI(β)) in the three conditions. Time course of average beta power modulation (15 – 30Hz) averaged over sensors of interest within the control group (blue) and ADHD group in the placebo (green) and MPH condition (red). Shaded error bar depicts standard error over subjects along timepoints. (B) Mean PI(β)s per condition, revealing a stronger beta suppression for the control group, compared to the ADHD group in the placebo condition (p=.016) and an increased beta suppression in the ADHD group when MPH was administered, as compared to placebo (p=.028). PI(β)s in the ADHD group on MPH didn’t differ significantly from the controls.

### Differences in beta preparation values prior to motor response are not attributable to differences in the baseline period

We next assessed whether the differences in PI(β)s across groups were due to different levels of beta power in the baseline period (i.e. ‘resting-state’ interval, prior to the onset of the cue). We hence computed raw power in the low frequencies (5-30Hz) locked to the onset of the cue (**Figure 7A**) and then selected 15 pairs of symmetric central sensors displaying highest raw beta (15-30Hz) power in the whole pre-cue interval (−500 – 0ms, cue-locked). The topographical representation of raw beta power in the selected interval averaged over groups is shown in **Figure 7B**. Mean beta power in each group (**Figure 7C**) was then compared between ADHD_MPH_ and ADHD_PLA_ by means of paired t-test (t_(17)_=.62, *p*=.540) and between TD and ADHD group in both conditions (ADHD_MPH_ vs TD: t_(42)_=.37, *p*=.711; ADHD_PLA_ vs TD: t_(42)_=−.20, *p*=.837) via independent sample t-test. Pairwise difference in raw beta power between the two ADHD conditions is depicted in **Figure 7D**. The results of this analysis suggests that the differences in PI(β)s previously observed were not explained by differences in raw beta power in the pre-cue interval (i.e. baseline time-window).

**Figure 7.**
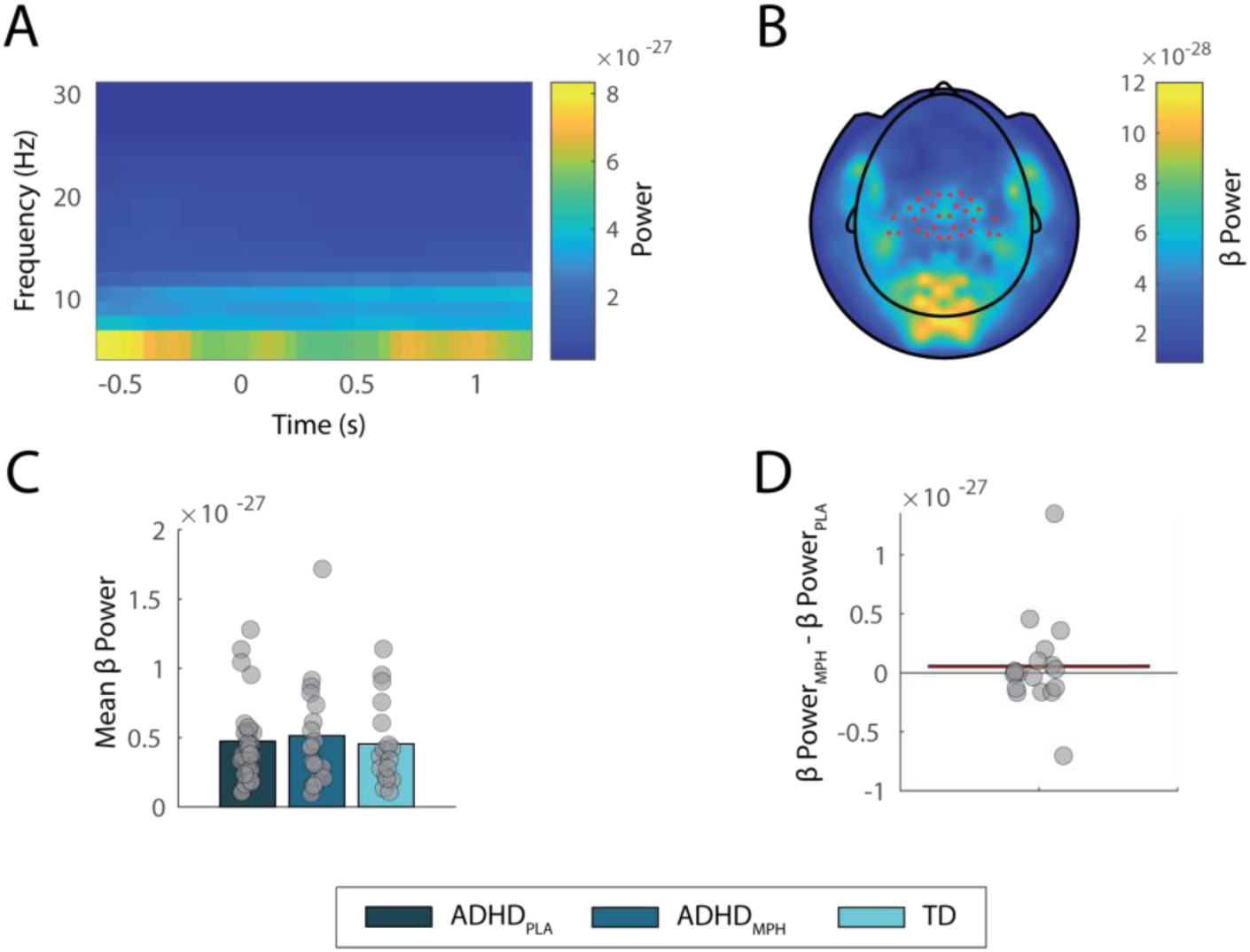
Raw beta power in the pre-cue interval in the three experimental groups. (A) Time frequency representation of raw power in the lower frequency bands, locked to the onset of the cue. (B) The selected pairs (N=15) of symmetrical central sensors displaying highest pre-cue beta power are highlighted in red in the topography, while average and single values per group are represented in panel C via bars and superimposed individual datapoints. No significant difference between ADHD and TD groups was identified, neither between conditions within the ADHD group. Panel D shows pairwise difference in raw beta power between ADHD_MPH_ and ADHD_PLA_.

### Behavioral correlates of beta desynchronization

To explore potential links between oscillatory and behavioral asymmetries, we examined the relationship between interhemispheric modulation of beta depression and perceptual and motor behavioral asymmetry, as indexed by individual LBT scores. To this purpose, we computed hemispheric modulation indices of beta modulation (HMI(β)) for each subject, separately for left and right ROIs (see *Materials and Methods*). The correlation between HMI(β) and LBT values was then considered in each experimental group. However, there was no significant correspondence between behavioral spatial biases and hemispheric beta modulation in either of the groups (ADHD_MPH_: r=.24, *p*=.327; ADHD_PLA_: r=.01, *p*=.940; TD: r=14, *p*=.491)

Based on previous research pointing to atypical beta preparation values in children diagnosed with ADHD, we next related symptoms severity (as reflected by parents’ rating on the ADHD scale) to the individual ability to suppress beta power in preparation to the motor response. We considered symptoms across both the TD and the ADHD (placebo condition) group, hence embracing the notion that ADHD symptomatology derives from a ‘spectrum’, rather than from a dichotomous distinction with TD peers. We then applied a general linear mixed model incorporating individual mean ADHD scores together with subjects’ age as regressors for the respective PI(β). Notably, we found that symptoms severity predicts beta depression during motor preparation, when controlling for age (full model R_2_=.27, *p=* .001; partial *p*=.017): specifically, the lower the symptoms, the stronger the beta preparation in the preparation interval (**Figure 8**).

**Figure 8.**
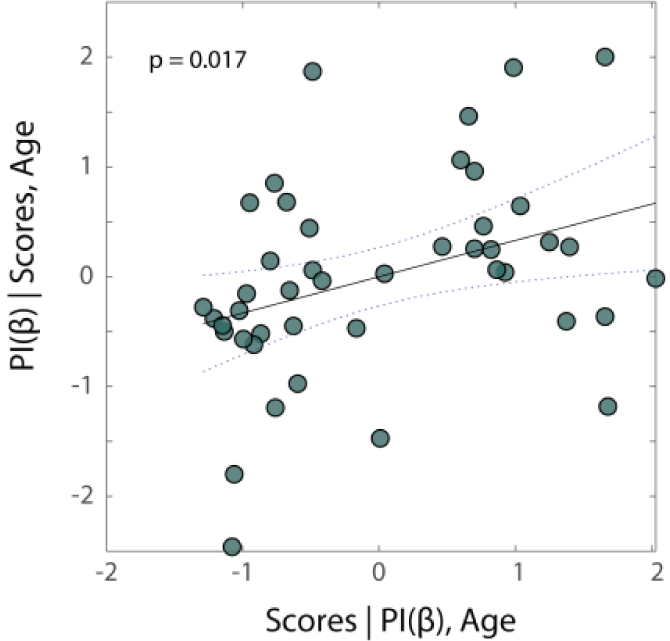
Severity of ADHD symptoms predicts beta preparation values in all subjects. Partial regression plot displaying the linear association between symptoms severity (‘Scores’) and beta preparation values PI(β) across all participants in the study (TD and ADHD (placebo) subjects merged in a single sample). A significant positive regression (p=.017) indicates that higher symptoms are predictive of lower beta preparation values in preparation to motor response to the cued target in the task.

### Beta depression predicts behavioural performance in the ADHD group in the MPH condition

To investigate the link between beta depression and task performance, we first correlated the beta preparation (PI(β)) with the inverse efficiency score (IES; i.e. median RTs divided by accuracy) over subjects (**Figure 9A**). No significant correlation was found when considering the data combined over groups (*p*=.274). When considering the three groups separately, a significant relationship emerged between PI(β) and behavioural performance in the ADHD_MPH_ group. To further evaluate this relationship, we specified a linear regression model, including the interaction PI(β)*Dosage for the prediction of IES values. A significant regression was found (R_2_=.58, *p*=.005), holding significant main effects (PI(β): *p*=.004, Dosage: *p*=.009) and interaction effects (PI(β)*Dosage: *p*=.006). **Figure 9B** shows that the interaction between beta depression and MPH dosage (PI(β)*Dosage) significantly explains behavioral performance (i.e. stronger beta depression predicts better performance).

**Figure 9.**
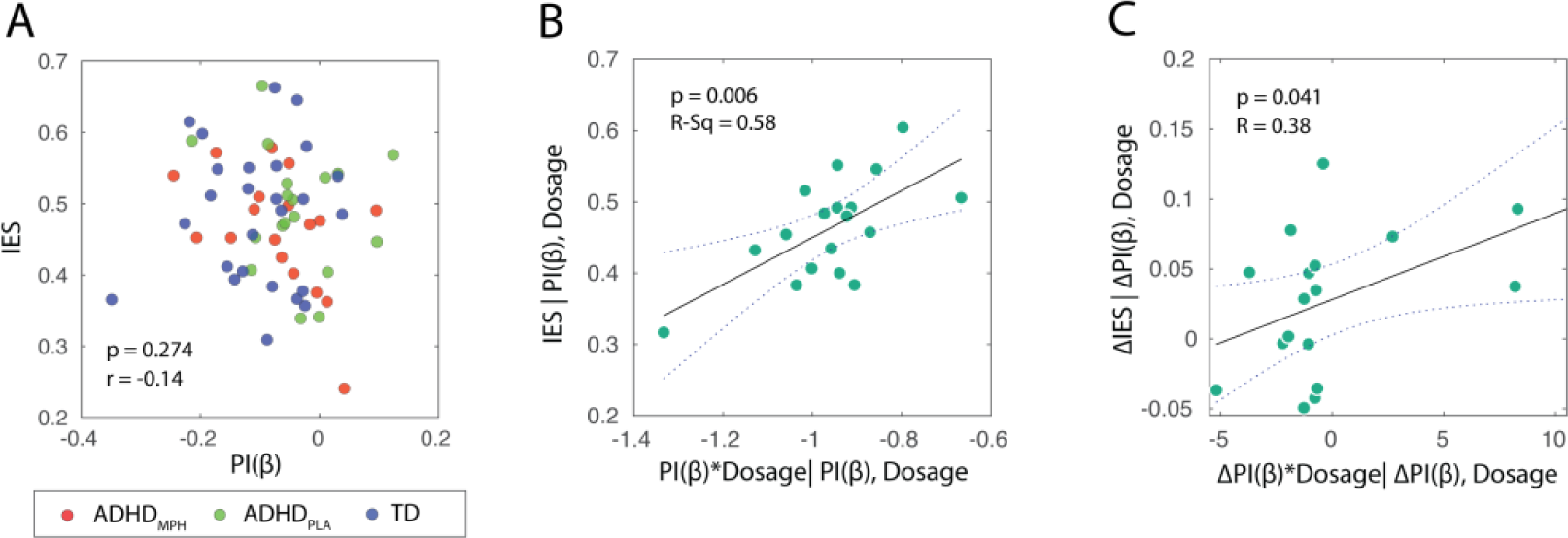
Interaction between medication dosage and beta preparation predicts task performance in ADHD_MPH_. (A) Association between IESs and PI(β)s at the overall subject population level, clustered in the scatter plot according to the categorical variable ‘group’. (B) Partial regression plot showing the association, in the ADHDMPH group, between IESs and PI(β)s, within a linear regression model controlling for dosage*PI(β) interaction (p=.006). (C) Partial regression plot depicting the association between Δβ-PI and ΔIES in the ADHD_MPH_ condition: when controlling for the effect of Dosage*Δβ-PI interaction, increase in beta preparation values due to medication intake (Δβ-PI) predicts improvements in behavioral performance (ΔIES) between MPH and placebo condition.

Similarly, we addressed whether the changes in beta depression due to medication could explain changes in behavioural performance in the task between placebo and MPH condition when controlling for the interaction with dosage administered. To determine the improvement in performance due to medication we computed the difference in IESs between placebo and MPH conditions as follows:

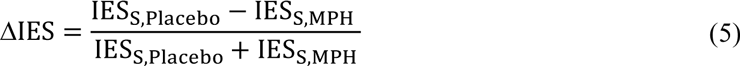

Consequently, a positive ΔIES for a given subject denoted better performance in the task when MPH was administered as compared to Placebo, and vice versa. We then quantified the difference in beta depression (PI(β)) between placebo and MPH conditions as:

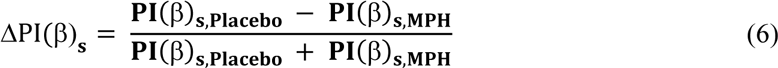

A positive ΔPI(β) change for a given subject *s* reflected stronger beta preparation index with MPH compared to placebo – and vice versa.

We hence employed a linear regression model analogous to the one previously described, in order to assess whether MPH induced changes in beta depression (ΔPI(β)) could explain improvements in behavioural performance in the task (ΔIES), including the interaction between ΔPI(β) and individual MPH dosage administered. A significant main effect for ΔPI(β) (*p*=.025) and a significant interaction ΔPI(β)*Dosage (*p*=.041) were found (**Figure 9C**).

### Beta power lateralization and hemispheric lateralized modulation biases

In order to assess interhemispheric biases in the modulation of beta power in preparation to the response, we computed beta lateralized modulation indices (LMI(β)) for each participant and for each experimental condition, according to Eq.1 (see *Materials and Methods*). Grand average LMI(β)s for left and right hemisphere ROIs are shown in **Figure 10**. Given that subjects were instructed to respond to left and right targets via button press with the left and right finger, respectively, we identified a clear lateralization of beta power, prior to the motor response, which was lower at central sensors contralateral to the target (i.e., contralateral to the hand used to respond).

We then investigated whether lateralized modulation of beta power (LMI(β)) differed between the three experimental groups (**Figure 10B, C, D**). For each subject within a group, we averaged LMI(β) over left and right ROIs (previously selected from the grand average). We first used a paired t-test between hemispheres, to evaluate whether there was a significant lateralization of power. **Figure 11A** shows mean LMI(β) values per hemisphere and in the three different groups confirming a significant lateralization as assessed by paired t-test between left and right hemisphere ROIs for ADHD_Placebo_ (t_(17)_=−3.39, *p*=.003), ADHD_MPH_ (t_(17)_=3.83, *p*=.001), and TD (t_(25)_=−4.03, *p*=4.5×10_-4_). LI(β)s were then derived for the three groups according to Eq(2), in order to assess whether significant differences occurred in beta lateralization due to medication intake and between groups. Mean LI(β) values for the three groups are depicted in **Figure 11B**. No significant differences in LI(β) were found within the ADHD group, when comparing MPH and Placebo condition (t_(17)_=1.03, *p*=.32), neither between TD and ADHD_Placebo_ (t_(42)_=.98, *p*=33) nor ADHD_MPH_ (t_(42)_=1.71, *p*=0.09).

**Figure 10.**
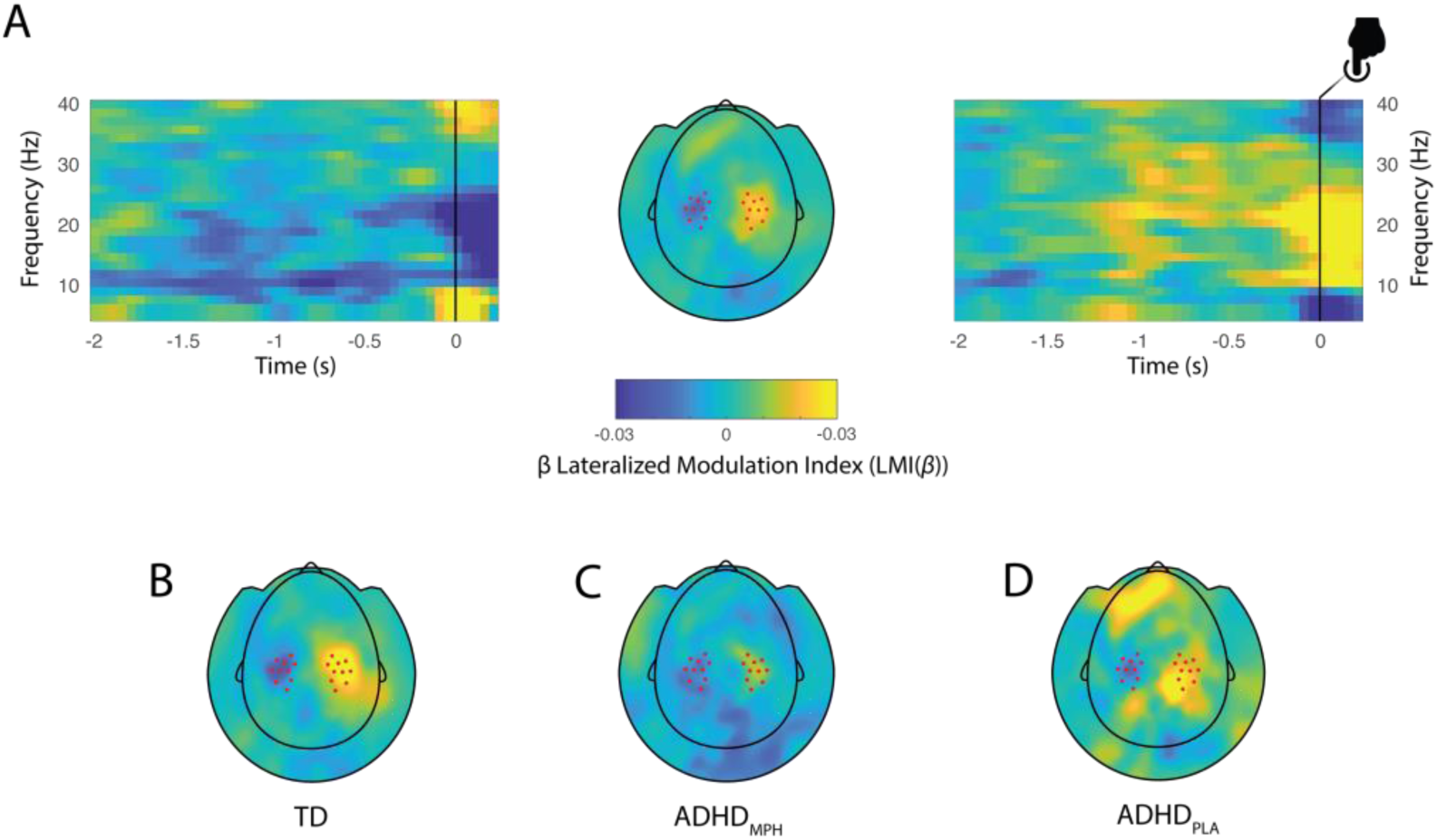
Grandaverage of hemispheric lateralization of beta modulation. (A) Time-frequency representations of power (TFRs) and topographical plot showing contrast between the ‘cued right’ – ‘cued left’ trials. A clear beta band lateralization is visible at central sensors in the in the −1000 – 0ms interval. Sensors included in the left and right ROIs are marked as dots in the central topoplot. Trials are locked to the onset of the motor response. On the bottom, topographic plots of beta lateralization are displayed for the TD (B), ADHD_MPH_ (C) and ADHD_Placebo_ (D) conditions. Red dots on the topographies highlight the cluster of sensors of interest considered for the computation of the TFRs, corresponding to the same symmetric pairs identified

**Figure 11.**
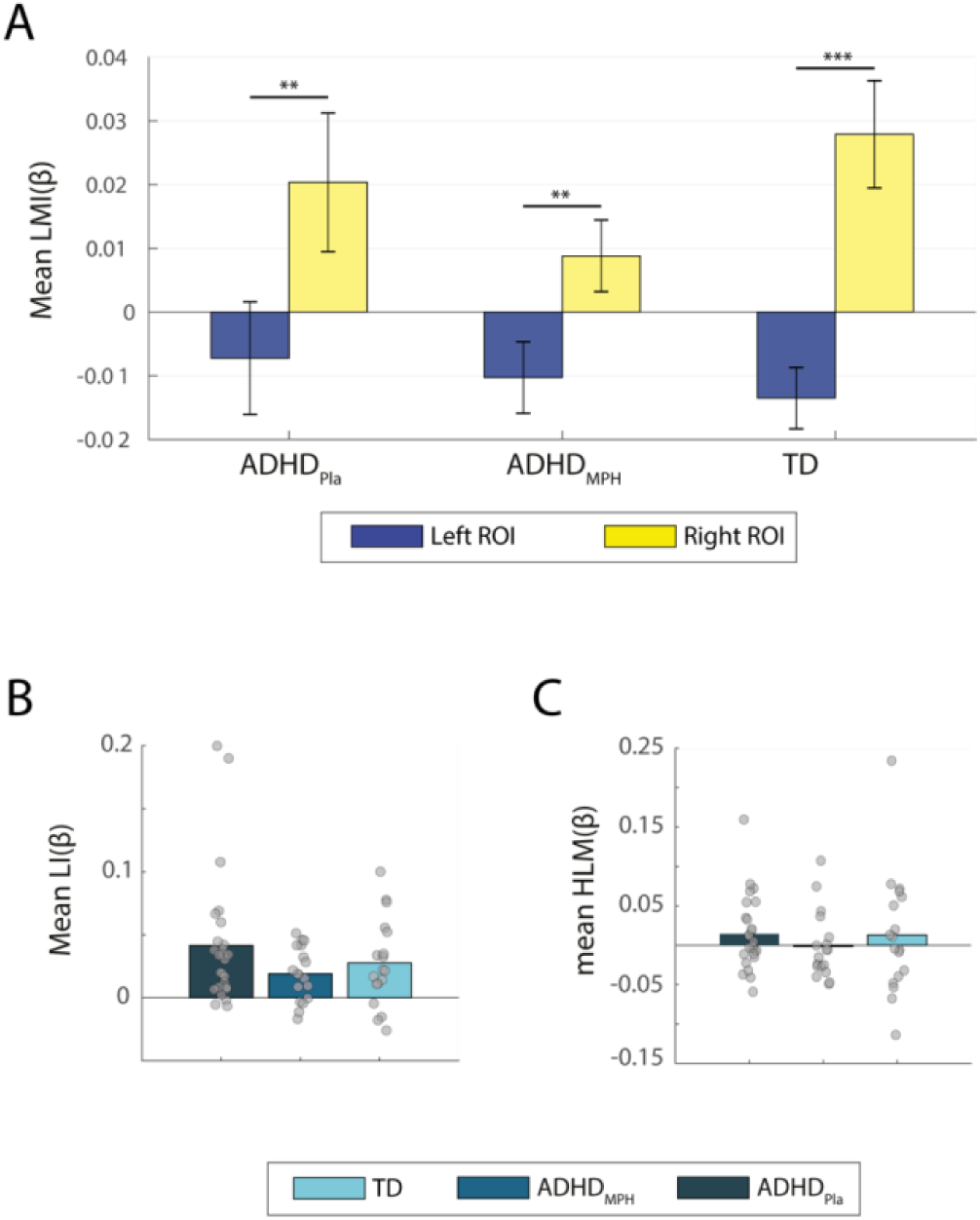
Modulation indices per hemisphere and related t-test per experimental condition. (A) Hemispheric beta lateralization for left (blue) and right (yellow) ROIs, for the three groups. Dependent samples t-test was used to assess difference in beta lateralization between hemispheres. (B) Mean beta lateralization indices for the three experimental conditions. No significant difference was found between groups as assessed by dependent (ADHD_MPH_ vs ADHD_Placebo_) or independent samples t-test (ADHD_MPH_ vs Controls, ADHD_Placebo_ vs Controls). Asterisks denote statistical significance. (C) Mean hemispheric lateralized modulation indices per experimental condition. These indices indicated whether, on average, there was a leftward (negative value) or rightward (positive value) bias in the ability to modulate beta power in the left or right hemisphere. None of the groups displayed this bias, as assessed by one sample t-test.

In order to assess potential hemispheric biases in the modulation of beta power, we computed indices of hemispheric lateralized modulation of beta (HLM(β)) per subject, according to Eq.(3). Mean HLM(β) indices per condition are presented in **Figure 11C**. Given that the assumption of normality was not met for the distribution of HLM(β)s across groups, we used two-sided Wilcoxon signed rank test to assess significant left or rightward biases. We found that none of the three groups displayed a significant hemispheric bias in lateralized beta modulation (ADHD_MPH_: *z* = −.76, *p*=.450; ADHD_Placebo_: *z* = .63, *p* =.530; TD: *z* = 1.13, *p* =.260).

### Behavioral correlates of beta lateralization

### Stronger beta lateralization is associated with better task performance in the TD group

To examine the association between the beta lateralization values (LI(β)), in preparation to motor response and task performance, we correlated LI(β) with CEI and IES in the three different conditions. **Figure 12** shows scatter plots of the linear association between CEI and LI(β) for the three experimental groups. No significant relationship was found between behavioural performance and beta lateralization in the ADHD_Placebo_ (CEI: r=−.46, *p*=.052; IES: r=−.15, *p*=.560), nor in the ADHD_MPH_ (CEI: r=.-23, p=.361; IES: r=.10, *p*=.708) conditions, whilst a significant negative correlation emerged between behavioral measures and LI(β) in the TD group (CEI: r=−.85, *p*=3.5×10_-8_; IES: r=−.49, *p*=.012). A comparison of independent correlation coefficients, according to the methods described in (Wilcox, 2009), confirmed that the association between behavioral performance (CEI) and beta lateralization (LI(β)) was significantly higher in the controls as compared to ADHD_MPH_ group (95% CI: [0.16–1.18]), although not reaching significance when compared to MPH_Placebo_ (95% CI: [−0.08–1.12]).

**Figure 12.**
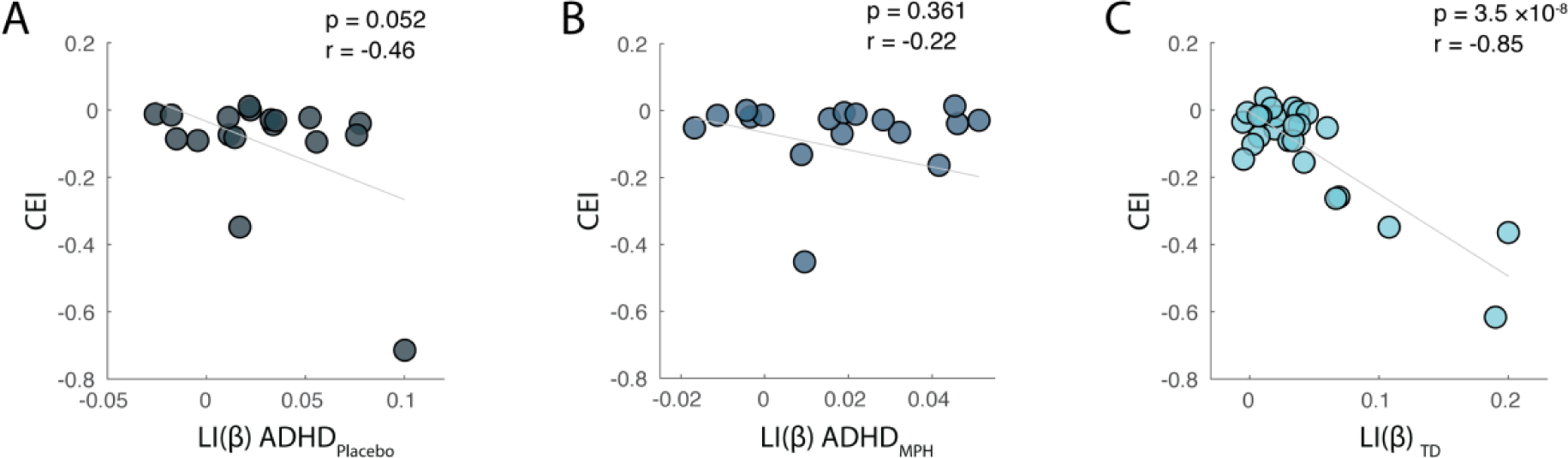
Association between Cueing Effect Index (CEI) and beta lateralization in the three experimental groups. Correlation between beta lateralization indices and behavioural performance expressed in terms of cue benefit (calculated on IES scores) in the three groups (A: ADHD_PLA_, B: ADHD_MPH_, C:TD). For the controls, a significant relationship was found: stronger beta power lateralization was associated with better behavioural performance in the task. A pairwise comparison between independent correlations showed a significant difference between ADHD_MPH_ and TD correlation coefficients between LI(β) and CEI.

To further confirm this association, we divided the TD group by the median split, according to the CEI. As a result, we obtained two subgroups of N=13 (namely ‘*high cue effect’* vs ‘*low cueing effect*’) and then averaged the TFRs of lateralized beta modulation indices (LMI(β)) associated with the two subgroups over the time-frequency spectrum of interest. **Figure 13** displays a topographical representation of LMI(β) values per subgroup (A, B), together with individual raw data points, superimposed on bars representing average values per ROIs per each subgroup (C, D).

**Figure 13.**
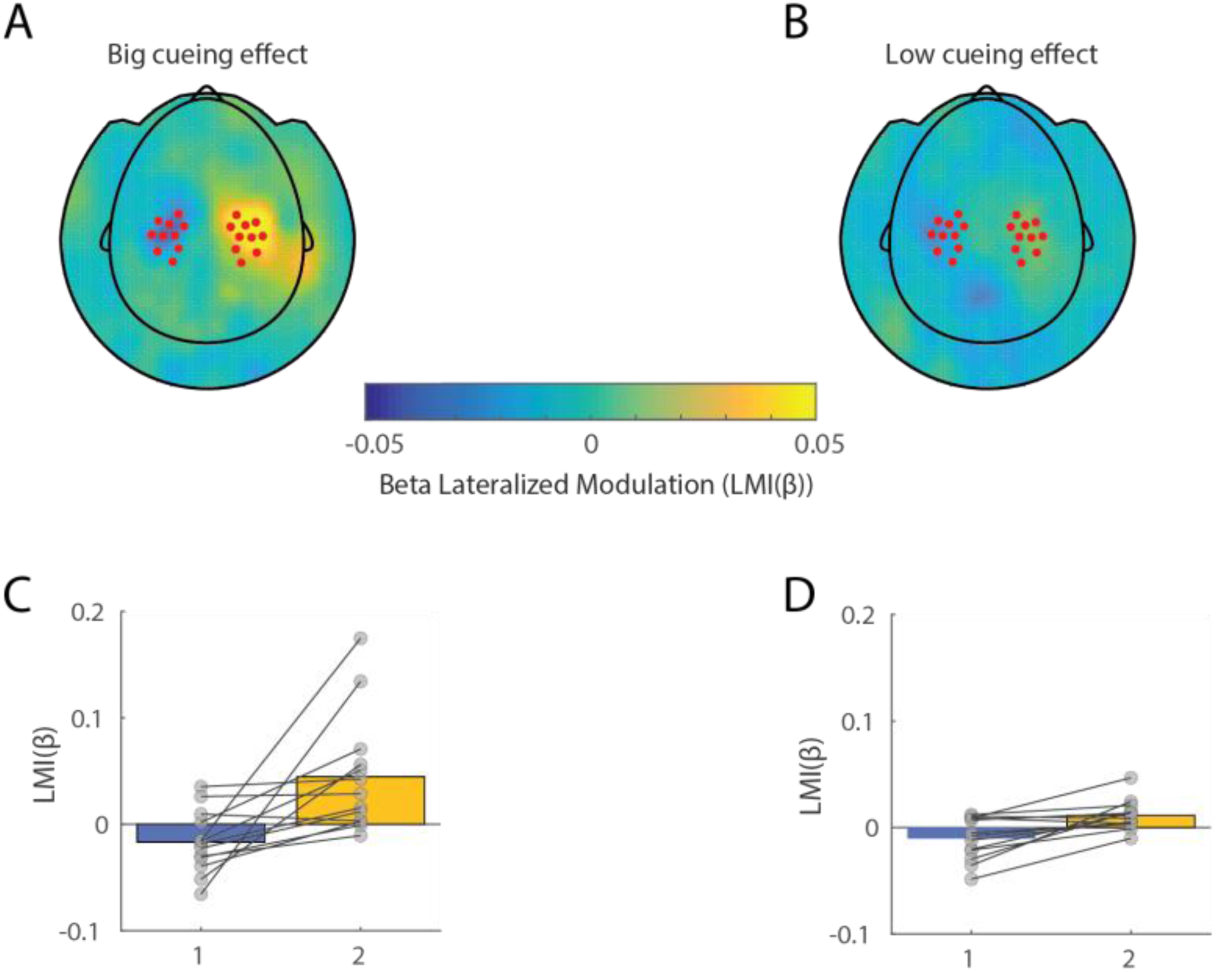
Beta lateralized modulation indices in relation to task performance. Topographical plot of LMI(β) values for two subgroups of the TD subjects, clustered according to degree of cue benefit (high (A) vs low (B) Cueing Effect Index (CEI)). Left and right sensors of interest are marked as dots and correspond to the same ROIs as in Figure 9. Individual datapoints are superimposed on bar graph showing individual scores and LMI(β) averaged over ROIs in the high (C) and low (D) CEI subgroup. Average LI(β) (see Eq.(2)) was stronger for participants who benefited more from the cue direction in terms of task performance, as compared to the ones displaying a lower CEI.

Next, we generated LI(β)s indices for *high* vs *low* cueing effect according to Eq.(2) and statistically assessed their difference by means of dependent sample t-test. This revealed that subjects who displayed a stronger beta lateralization in preparation to motor responses to a cued target, also benefited more from the cue (t_(12)_=−2.15, *p*=0.049); however, this effect was absent in the ADHD group.

### Lateralized beta modulation and line bisection task

Analogously to the premises aforementioned for the beta depression analysis, we sought to identify, if any, the relationship between asymmetries in lateralized oscillatory activity in the beta band (HLM(β)) and perceptual and motor asymmetries reflected by performance in the LBT. We hence correlated HLM(β) values for each group with the respective score in the LBT task. As a result, no significant association was found between the two indices for neither of the three experimental groups (ADHD_MPH_: r=−.06, *p*=.798; ADHD_Placebo_: r=−.024, *p*=.321; TD: r=.01, *p*=.924)

### Alpha modulation decreases with MPH administration in the ADHD group

To quantify patterns of alpha modulatory activity in the preparation interval following the cue, we first baseline-corrected TFRs for all subjects with respect to 500ms prior the onset of the cue. Baselined were then averaged in the three experimental groups, with the aim to select a common set of sensors displaying the strongest modulation in the alpha band, to be used in further analyses. To this end, we considered 40 pairs of symmetrical sensors belonging to the occipital and parietal areas, displaying highest alpha power (*f*=7 – 13Hz) in a time window 400 – 1200ms post-cue (excluding early evoked activity). Time-frequency and topographical representation of power (baseline corrected) were then computed for each group separately, by averaging over the sensors (**Figure 14 A, B, C**, right panel) and the TFRs (**Figure 15 A, B, C**, left panel) of interest. Average alpha modulation indices for these sensors and in the TFR, were then computed for each group separately (**Figure 14D**) and referred to as alpha retention index (RI(α)), while a temporal evolution of their value is depicted in **Figure 14E**.

**Figure 14.**
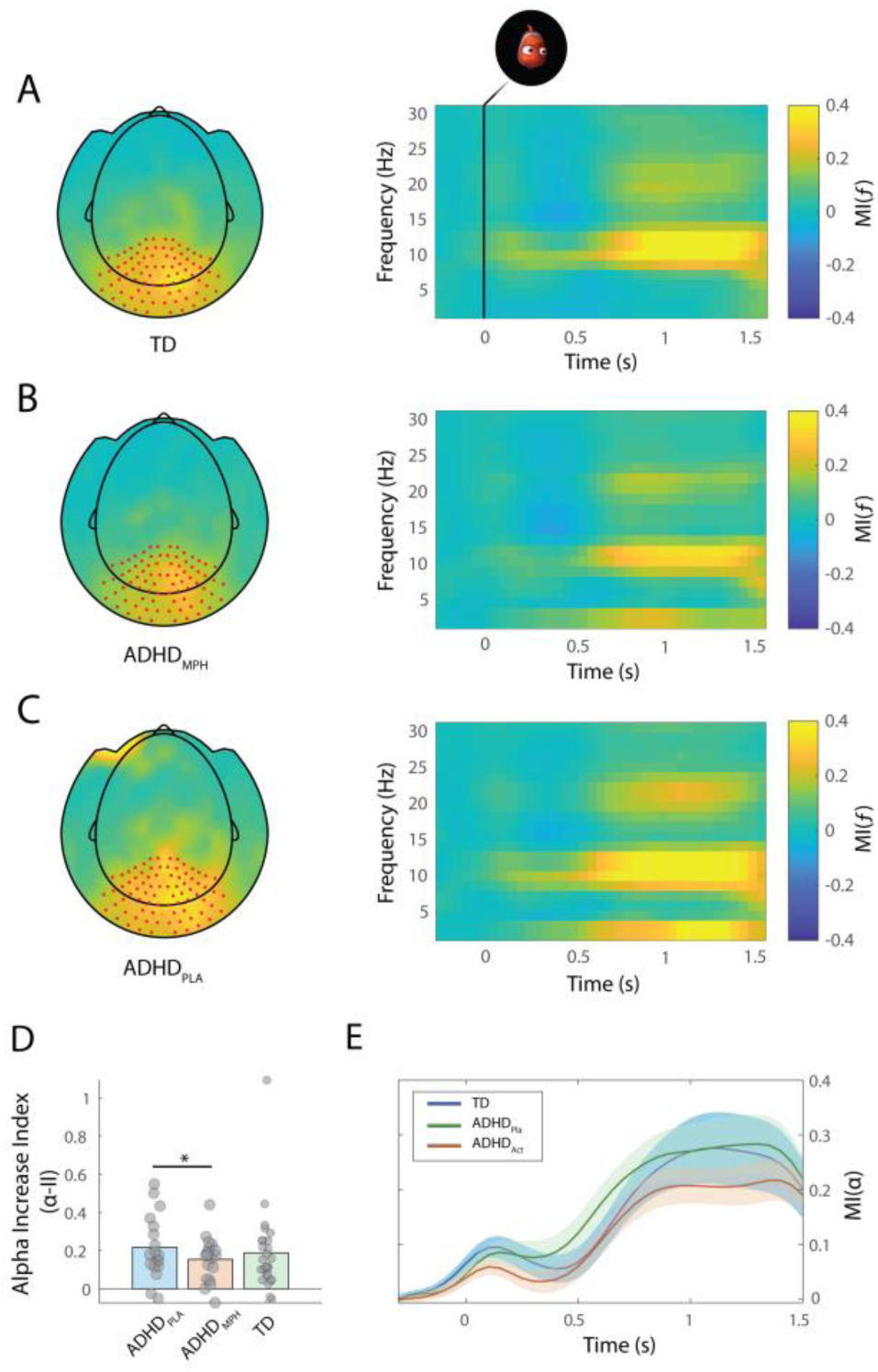
Alpha power modulation in three groups Time course of alpha power increase and mean power increase per condition. Topographic plots (left) and respective time-frequency representation of power modulation following cue onset in the TD (A), ADHD_MPH_ (B) and ADHD_Pla_ (C) group. A clear alpha power increase is visible at posterior sensors which is reduced in the ADHD group in the MPH condition (*p*=.048). (D) Mean RI(α) values per condition; the individual subject data are superimposed on the bars. (E) Temporal evolution of MI(α) locked to the onset of the cue (t=0), along the time window for the three experimental groups.

**Figure 15.**
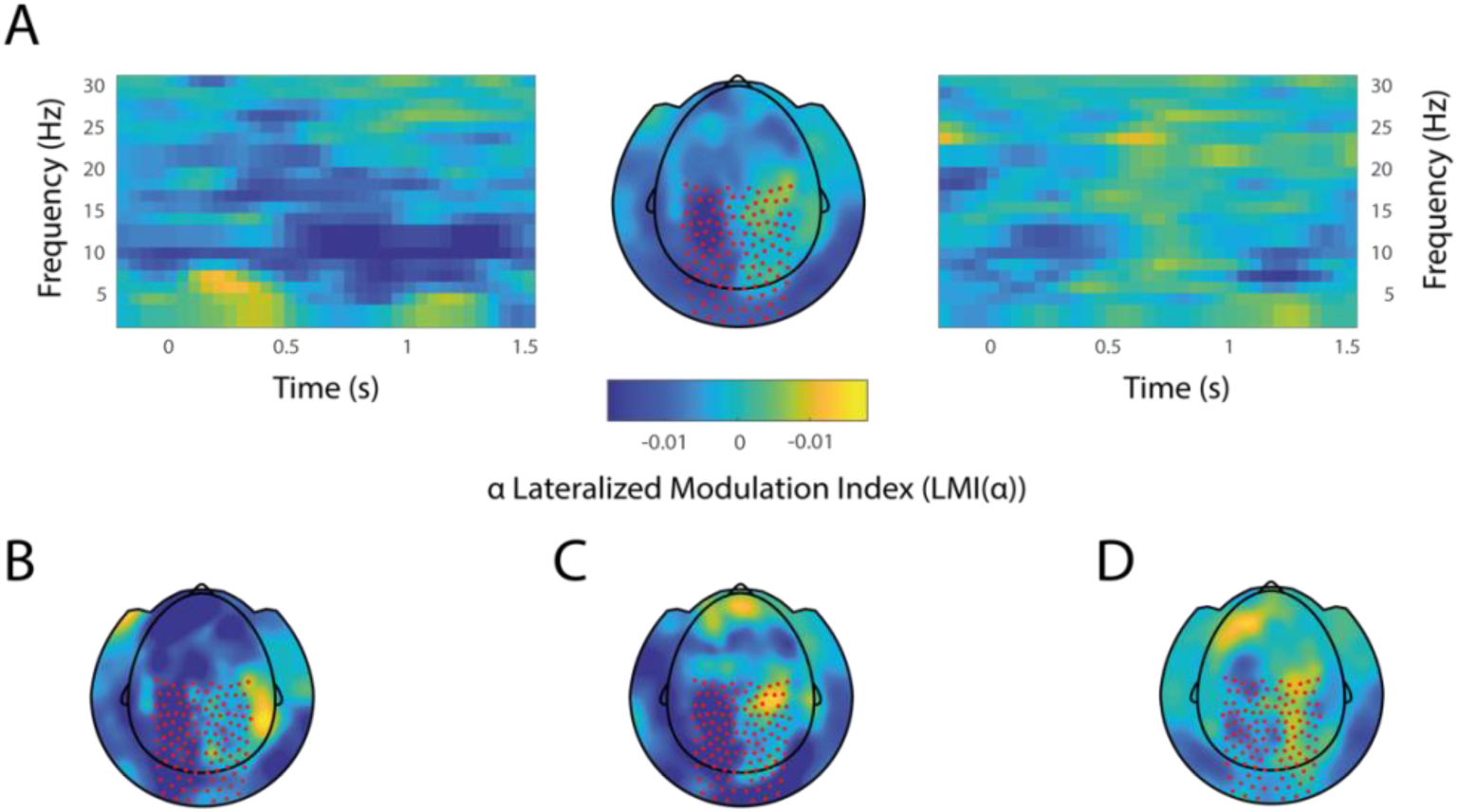
Alpha power lateralization in the three conditions. (A) Left and right-side panels show the grand average time-frequency representation of lateralized alpha power following cue onset (LMI(α)) averaged over the left and right ROIs, respectively. Central panel represents the topographical representation of LMI(α) averaged over time and frequency spectrum of interest. Sensors of interest are market as red dots on the topography. Topographical representations of grand average LMI(α) for the TD(B), ADHD_MPH_ (C) and ADHD_Pla_ (D) group. When evaluating the significance of alpha lateralization based on interhemispheric power difference comparison via paired t-test, only the ADHD_Pla_ group showed a significant lateralization of power (*p*=.014).

A clear increase in alpha modulation was present across the cue-target interval during retention of cue information. We hence quantified the degree of alpha power retention (RI(α)) separately in the three groups by averaging the MI(α) over the cluster of sensors selected. An independent sample t-test was employed to assess statistical difference between RI(α) in the TD group and in the ADHD group for both conditions, while a paired t-test evaluated the comparison between MI(α)_MPH_ and MI(α)_PLA_. Importantly, we found a significant difference between conditions in the ADHD group (t_(17)_=2.12, *p*=.048), indicating that MPH administration led to a significant decrease in alpha power retention (RI(α)).No significant difference in alpha retention arose instead between TD and ADHD_PLA_ (t_(42)_=−.46, *p*=.642) nor between TD and ADHD_MPH_ (t_(42)_=.62, *p*=.537),

### Differences in alpha retention in the preparation interval are not attributable to differences in the baseline period

According to the same methods applied in the analysis of beta, we ensured that the differences in RI(α)s across groups were not due to different levels of alpha power in the baseline period. We then considered the raw low frequency power prior to the cue, and selected 20 pairs of sensors showing highest alpha retention in the 500ms prior to the onset of the cue. Mean alpha power in each group was then compared between ADHD_MPH_ and ADHD_PLA_ by means of paired t-test (t_(17)_=.95, *p*=.355) and between TD and ADHD group in both conditions (ADHD_MPH_ vs TD: t_(42)_=1.15, *p*=.254; ADHD_PLA_ vs TD: t_(42)_=.69, *p*=.497) via independent sample t-test. Pairwise difference in raw alpha power between the two ADHD conditions is depicted in **Figure 7D**. The results of this analysis suggest that the differences in alpha power retention (RI(α)) previously observed were not explained by differences in raw alpha power in the pre-cue interval.

### Behavioral correlates of alpha retention

To explore potential links between oscillatory and behavioral hemispheric asymmetries, we examined the relationship between interhemispheric modulation of alpha retention and perceptual and motor behavioral asymmetry, as indexed by individual LBT scores. Following the same methods as for the beta range analysis, we derived hemispheric modulation indices of alpha modulation (HMI(α)) for each subject, separately for left and right ROIs (see *Materials and Methods*). Then, we computed Pearson correlations between HMI(α) and LBT values for each group. However, there was no significant relationship between behavioral spatial biases and hemispheric beta modulation in either of the groups (ADHD_MPH_: r=−.11, *p*=.663; ADHD_PLA_: r=−.25, *p*=.303; TD: r=−.22, *p*=.280)

### Alpha increase and behavioral performance

To assess the existence of an association between alpha increase and individual task performance, we first considered the correlation between alpha RI(α) and IES over subjects. No significant correlation was found when considering the data combined over groups (*p*=.450) nor when considering the three groups (ADHD_MPH_: r=−.22, *p*=.389; ADHD_PLA_: r=−.01, *p*=.970; TD: r=−.14, *p*=.502). For ADHD group on MPH, we employed a linear regression model, including the interaction RI(α)*Dosage for the prediction of IES values, which did not lead to significant main effects (RI(α): t=.69, *p*=.500; Dosage: t=1.19, *p*=.260) not interaction (t=−.78, *p*=.452).

Similarly, we addressed whether the reduced alpha increase due to medication could explain drug related changes in behavior (when controlling for dosage administered). To determine differences in RI(α) due to medication we computed ΔRI(α) according to:

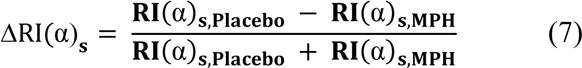

A positive ΔPI(α) change for a given subject *s* reflected stronger alpha increase in the placebo condition compared to MPH – and vice versa. We hence employed a linear regression model analogous to the one described for ΔPI(β), to assess whether MPH induced changes in alpha retention (ΔRI(α)) and the relation to MPH dosage, could explain improvements in behavioural performance in the task (ΔIES). No significant main effects (ΔRI(α): t=−.04, *p*=.970; Dosage: t=−.63, *p*=.541) nor interactions (t=.03, *p* =.971) were found.

### Alpha power lateralization

In order to assess interhemispheric biases in the modulation of alpha power following the cue, we computed alpha lateralized modulation indices (LMI(α)) for each participant and for each experimental condition, according to Eq.1, with *f=* 7-13Hz (see *Materials and Methods*). Grand average LMI(α)s for left and right hemisphere ROIs are shown in **Figure 13**. A lateralization of alpha power was visible at posterior (parieto-occipital) sensors, showing a lower LMI(α) in the left ROI (contralateral to the target (i.e., cue direction)). We then investigated whether lateralized modulation of alpha power (LMI(α)) differed between the three experimental groups. For each subject within a group, we averaged LMI(α) over left and right ROIs (previously selected from the grand average). We first used a paired t-test between hemispheres, to evaluate whether there was a significant lateralization of power. While no significant interhemispheric difference in alpha lateralized power was found for the TD and the ADHD_MPH_ (t_(25)_=−1.95, *p*=.06 and t_(17)_= −1.75, *p*=.09), the ADHD_PLA_ group displayed a significant interhemispheric LMI(α) (t_(17)_=−2.73, *p*=.014). LI(α)s were then derived for the three groups according to Eq (2), in order to assess whether significant differences occurred in beta lateralization due to medication. No significant differences in LI(α) were found in the ADHD group, when comparing MPH and Placebo condition (t_(17)_=.52, *p*=.610); neither between TD and ADHD_Placebo_ (t_(42)_=1.20, *p*=.236) nor ADHD_MPH_ (t_(42)_=0.62, *p*= .540).

In order to assess potential hemispheric biases in the modulation of alpha power, we computed indices of hemispheric lateralized modulation of alpha (HLM(α)) per subject, according to Eq.(3). No hemispheric bias in HLM(α) were found when evaluating, by means of t-test, whether any significant left or rightward biases in lateralized modulation existed in the three groups (ADHD_MPH_: t_(17)_=−1.71, *p*= .106; ADHD_Placebo_: t_(17)_=−1.96, *p*= .065; TD: t_(25)_=−0.12, *p*= .905).

### Behavioral correlates of alpha lateralization

Given the premise that a significant interaction existed between *session order* (first vs second session) and *drug* (MPH vs Pla) in the estimation of LI(α) values (see **Table 3**), we employed a general linear model, including the abovementioned interaction, to investigate whether LI(α) values could predict behavioral performance. The model included all two-way interactions, excluding the three-way interaction LI(α)\**drug*\**session*. The model was not overall significant (R_2_ = −.07, *p*=.716) and no significant main effects nor interaction were found for the terms.

As for beta lateralization, we also assessed a potential relationship between asymmetries in lateralized oscillatory activity in the alpha band (HLM(α)) and perceptual and motor asymmetries reflected by performance in the LBT. Following the same logic above, we built a general linear model for the prediction of LBT values, including HLM(α), *drug* and *session order* and their interaction as regressors. Also in this case, the model specified could not significantly predict LBT values (R_2_ = −.11, *p*=.878).

## Discussion

We compared somatosensory and attentional-related oscillatory dynamics reflecting motor preparation in children with ADHD and TD controls. Next, we implemented a double-blind, placebo-controlled crossover design to investigate the effects of stimulant medication (MPH) on these oscillatory patterns in children with ADHD.

### Beta oscillations

As expected, we first observed an overall beta band modulation preceding the onset of the motor response, expressed by a beta power depression (PI(β)). We show that mean PI(β) was stronger in the typically developing (TD) group as compared to the ADHD_Placebo_ group. Importantly MPH significantly increased the beta depression in the ADHD_MPH_ group, such that it was comparable to healthy controls. We also showed that the interaction between PI(β) and administered dosage of MPH, significantly explained performance in the task in the ADHD group: subjects whose beta depression increased more with MPH also were the subjects whose behavioural performance improved in the MPH as compared to the placebo condition. These interhemispheric beta oscillatory patterns in somatosensory cortex have been indeed consistently related with appropriate motor planning in healthy subjects and found to be altered in pathologies associated with movement disorders (Proudfoot et al., 2017).

We also quantified hemispheric beta power lateralization (LI(β)) with respect to the directional cueing (hence the hand used to respond) in the three groups, which is indicative of overall interhemispheric modulation of beta power prior to response to the target. Although we did not find a significant effect of MPH on beta lateralization, we did demonstrate that LI(β) is predictive of behavioural performance in the TD group.

Our findings suggest that MPH restores aberrant sensorimotor beta oscillations in children affected by ADHD, bringing the modulation closer to the one observed in the typically developing children. To the best of our knowledge, this is the first MEG study showing task related changes due to MPH in sensorimotor beta power in children with ADHD. A significant number of investigations have indeed focused on identifying different electrophysiological biomarkers of ADHD. A substantial amount of EEG research has been relating increased theta/beta ratio to the disorder (Ogrim et al., 2012; Gloss et al., 2016), although the robustness of these findings have been questioned by recent meta-analyses (Arns et al., 2013; Lenartowicz et al., 2018).

Fairly recent investigations have already shown aberrant sensorimotor activity, as indexed by mu oscillations, in relation to motor preparation, in individuals with ADHD (Yordanova et al., 2013; ter Huurne et al., 2018). These studies, in particular, suggest how improper inhibition and activation of motor cortical responses may be responsible for a suboptimal performance in children with ADHD. Our results corroborate the role of preparatory sensorimotor oscillations as electrophysiological correlates of the disorder, and further strengthen this link by showing the efficacy of MPH in normalizing this pattern.

Beta power decreases prior to a response to a cued location is known to index motor preparation (Tzagarakis et al., 2015). It hence is likely that behavioural motor symptoms in children with ADHD are reflected by a reduced ability to suppress sensorimotor beta power in tasks demanding readiness to a cued target. Interestingly, MPH acts by reinstating ‘regular’ beta modulation, providing indications for its mechanisms of action in the brain.

Which brain networks are modulated by MPH administration? MPH acts as dopamine (DA) agonist, thus enhancing dopaminergic availability in the synaptic cleft by inhibiting the DA transporter protein at the level of the striatum (Kodama et al., 2017). Striatal dysfunction in ADHD has indeed been documented (Rubia et al., 2011), and recently extended into a broader perspective of cortico-striatal-thalamic dysregulation (Mehler-Wex et al., 2006; Jenkinson and Brown, 2011; Peters et al., 2016).

Importantly, aberrant patterns of cortical beta activations have been observed also in Parkinson’s disease (PD) (Jenkinson and Brown, 2011), a neurodegenerative disorder known to be related to the degeneration of striatal dopaminergic neurons (Dostrovsky et al., 2002). Similarly to the effects of MPH reported in the context of the current study, direct evidence shows that both pharmacological therapy with L-dopa and subthalamic deep brain stimulation are able to normalize beta modulation, in PD patients, concomitant to their clinical improvement (Silberstein et al., 2005; Hammond et al., 2007). Our results corroborate the hypothesized causal link between striatal dopamine depletion and anomalous beta oscillations and provides an interesting conceptual bridge between the symptomatology of PD and ADHD patients, which partially share a common neural basis.

In ADHD research, a stronger focus has been mostly applied to the study of aberrant cognitive functions, which are then ‘restored’ by the use of MPH, from attention to working memory (Advokat and Scheithauer, 2013; Linssen et al., 2014; Ter Huurne et al., 2015; Fallon et al., 2016). These effects can be associated to reduced inattentive symptoms in ADHD patients treated with stimulant medication, coupled with their behavioural improvement. On the other hand, the hyperkinetic-impulsive behavior is an integral part of the disorder, to be considered in conjunction with attentional symptoms, particularly when considering the combined-type expression of ADHD. Importantly, it has been suggested that the interaction between cognitive and motor functions, rather than their single engagement, might be at the basis of aberrant stimulus processing in ADHD and the main functional target of stimulant medication (Berger et al., 2018). In our sample, MPH intake led to improved task performance and was reflected by the normalization of beta depression values in ADHD children.

In this regard, it is interesting to note that the commonality previously mentioned between PD and ADHD extends also to the attentional domain: in PD, enhanced bottom-up influence occurs to the detriment of top–down control of visual attention, which has been hence linked to striatal deterioration (Tommasi et al., 2015).

Altogether, prior work and our findings support the notion that beta band oscillations are reflecting not only motor planning/preparation, but flexible cognitive control and relevant attentional allocation at early stages of information processing (Engel and Fries, 2010; Hale et al., 2010; Haegens et al., 2017). Within this framework, beta-band oscillations in somatosensory cortex during motor preparation to a cued-target, might be involved in the top-down control mechanisms involved in attentional selection (Richter et al., 2018).

### Alpha oscillations

In the context of ADHD research, promising studies have been focused on alpha oscillations, in relation to both attention (Mazaheri et al., 2014; Vollebregt et al., 2015, 2016; Arns et al., 2018) working memory (Lenartowicz et al., 2016).

In the current study, we first observed a significant increase in posterior alpha power in all subjects (both groups and both conditions within ADHD group) preforming the attentional task, following cue onset, underlying the retention of cue information. Alpha power increases in visual cortex are linked to increased working memory (WM) allocation (Jensen, 2002; Sauseng et al., 2009; Jensen et al., 2012), and alpha power increases during vigilance tests (Loo et al., 2004) have been previously reported in children. Such increases in alpha power are believed to reflect reduced sensory excitability which in turn makes distractors less interfering, thus protecting working memory (WM) maintenance (Jensen, 2002; Haegens et al., 2010). Alpha oscillations during WM hence reflect functional inhibition of dorsal visual stream activity (Jokisch and Jensen, 2007), which enables stimulus processing in relevant cortical areas (e.g., motor preparation reflected by beta desynchronization). A corollary of this feature is the link between WM performance and alpha power increase (Haegens et al., 2010). In our sample, MPH led to an improvement in behavioral task performance as compared to the placebo condition, which, according to this established link, should then be reflected by a stronger alpha increase. Subjects’ abilities in WM tasks are negatively correlated with the ability to suppress distractors’ influence, hence, working memory load. This means that individuals with stronger WM capabilities are allegedly also the ones less affected by environmentally distracting input, needing less support from brain circuits involved in providing selective functional inhibition.

Consistently, recent studies revealed that different frequency bands underly event related synchronization during attention (10Hz) as compared to event related synchronization during WM retention (12Hz) and, in the case of retention, alpha peak magnitude increased with increased working memory load (Wianda and Ross, 2019). This finding is confirmed in the context of verbal WM studies, describing a negative correlation between alpha power modulation by memory load and WM capacity (defined as the ‘ability to sustain attention to the stored information in the face of interference or distraction’) (Hu et al., 2019; Proskovec et al., 2019).

Importantly, a previous EEG study has shown higher alpha power during a WM task in ADHD children compared to TD peers, reflecting weakened encoding capacity (Lenartowicz et al., 2019). This led to the conclusion that individuals with more efficient executive attention functions are also the ones less relying of sensory gating, reflected by alpha power increases. Consistently, given the beneficial effects of stimulants on executive functioning, among which attention and WM, children in the ADHD group administered with MPH also had to rely less on functional inhibition in order to retain information about upcoming target locations. These enhanced WM abilities were reflected by a weaker alpha power increase as compared to the placebo condition.

Our study offers important insights on the interplay of different neural dynamics that underpin ADHD and lie at the basis of the behavioral improvements observed with stimulant treatments. However, in order to identify the neural substrates of ADHD, a larger cohort must be considered. Such dataset would have the potential to assess the broader network involved in the generation of symptoms while properly controlling for their variability, given that unique structural and functional abnormalities are thought to be at the basis of the different subtypes (Lei et al., 2014; Mazaheri et al., 2014)

Future studies aimed at investigating the association between MPH induced beta oscillatory changes with individual striatal network properties would help to further elucidate the mechanisms of action of stimulant medication and provide more evidence on the role of beta oscillations and their relation to ADHD.

## Acknowledgements

The authors gratefully acknowledge the support of the Marie-Curie ITN grant ChildBrain (grant number 641652), the James S. McDonnell Foundation Understanding Human Cognition Collaborative Award (grant number 220020448), the Wellcome Trust Investigator Award in Science (grant number 207550) as well as the Royal Society Wolfson Research Merit Award.

